# A novel combination of CDK4/6 and PI3K inhibitors exhibits highly synergistic activity and translational potential in Ewing sarcoma

**DOI:** 10.1101/2024.10.14.618327

**Authors:** Maria Ana Isabel C. De Los Santos, Helen F. Gloege, Min Shen, Caleb O. Heaslip, Alejandra Aguilar-Quintero, Brigit McDannel, Mohammad Tanhaemami, Stephanie Meyer, Kelly Klega, Ya-Qin Zhang, David S. Shulman, Steven G. DuBois, Matthew Hall, Anne-Florence Blandin, Brian D. Crompton

## Abstract

Ewing sarcoma is a highly aggressive solid malignancy affecting children and young adults. Ewing sarcoma is driven primarily by EWSR1::FLI1, a fusion oncoprotein that has been notoriously difficult to target with traditional pharmacologic agents. There are numerous examples of preclinical promising combinations of small molecules that are never tested in pediatric clinical trials because agents fail to reach the market due to limited efficacy for common adult cancers. Moreover, the effectiveness of single-agent therapies for cancer treatment is often limited. To address these limitations, we selected 28 compounds that were largely FDA approved or in late stages of clinical development and known to regulate important pathways in Ewing sarcoma. We performed a drug screen in Ewing sarcoma cell lines with 180 combinations of tyrosine kinase inhibitors, cell cycle inhibitors, and conventional chemotherapy. The results of the screen revealed that a PI3K inhibitor, copanlisib, combined with a CDK4/6 inhibitor, ribociclib, exhibited strong synergistic anti-Ewing sarcoma activity. Using proteomic methods such as a reverse-phase protein array and western immunoblotting, we demonstrated that this combination induced a downregulation of the PI3K/AKT pathway as well as proteins involved in cell cycle regulation. We further confirmed these *in vitro* data using bulk RNA-sequencing. To evaluate the phenotypic effect of the PI3K/CDK4/6 inhibition in Ewing sarcoma lines, we performed apoptosis and cell cycle analyses using flow cytometry and demonstrated that ribociclib primarily induced a G0/G1 arrest with minimal effect on Ewing cell viability but significantly enhanced the apoptotic effect of copanlisib treatment. In a xenograft model of Ewing sarcoma, we demonstrated that the combination therapy significantly prolonged survival compared to treatment with either vehicle or single-agent therapy alone. Our findings identify a new candidate therapy combination for Ewing sarcoma using FDA-approved drugs and provide a resource of additional potential synergistic combinations for future validation.

**Statement of translational relevance:** Treatment of patients with newly diagnosed Ewing sarcoma involves intensive multi-agent chemotherapy, radiation, and surgery, often leading to long-term toxicities. Children with relapsed or metastatic disease experience poor outcomes with five-year overall survival rates of only 15 to 30 percent. Therefore, new therapeutic approaches are urgently needed. Our drug screen revealed several promising combinations with synergistic anti-Ewing sarcoma activity including the PI3K inhibitor copanlisib with the CDK4/6 inhibitor ribociclib. This combination also significantly improved survival in a mouse xenograft model of Ewing sarcoma compared to treatment with vehicle or either drug alone. Pre-clinical validation of a combination therapy composed of two FDA-approved drugs nominates this combination for early phase trials for patients with with relapsed or refractory Ewing sarcoma, a group of patients greatly in need of new therapeutic opportunities. The data generated by our drug screen may also be used as a resource to identify other promising combinations for future validation as Ewing sarcoma therapies.

## Introduction

Ewing sarcoma is an aggressive pediatric malignancy of bone and soft tissue affecting children, adolescents, and young adults. Treatment approaches for newly diagnosed patients consist of multi-agent adjuvant chemotherapy. Despite 5-year survival rates approaching 75% for those with localized disease, therapies have been largely ineffective for treating patients with metastatic or recurrent diseases (1–4). In addition, curative chemotherapy imparts considerable long-term toxicity and morbidity, affecting quality of life for survivors (5). Therefore, it is essential to find new therapeutic strategies for these young patients to reduce these adverse effects and improve overall outcomes.

Compared to adult cancers, pediatric cancers, particularly Ewing sarcoma, are characterized by a quiet genome with a low tumor mutational burden (6–13). Indeed, the hallmark molecular aberration identified in Ewing sarcoma is the presence of a reciprocal translocation involving FET family proteins and the ETS family of transcription factors, with *EWSR1::FLI1* and *EWSR1::ERG* translocations being the most common. Although Ewing sarcoma cells have a strict dependency on these fusion proteins, they act as pioneer transcription factors that lack a defined enzymatic pocket for ligand interaction making them a challenging pharmacological target (14).

In the absence of an immediately available compound or therapy that directly suppresses EWSR1::FLI1 activity, we chose to focus on combinations that target signaling pathways that are known to be dependencies in Ewing sarcoma. In previous work, we and others have identified dependencies on mTOR pathway activity and cell-cycle regulation (15–24). Targeting these pathways with small molecular inhibitors of Focal Adhesion Kinase (FAK) and Aurora Kinase B (AURKB), for example, has demonstrated synergistic inhibition of cell viability in Ewing sarcoma cells (25). Despite these encouraging results, access to these inhibitors for clinical trials has remained elusive.

To address the immediate need for therapeutic combinations for patients with Ewing sarcoma, we undertook a focused *in vitro* drug screen designed to identify commercially available drugs with synergistic anti-Ewing activity. For this screen, we purposefully selected agents that were either FDA-approved cancer drugs or were in late-stage clinical development at the time of the screen to expedite the translation of promising combinations into clinical trials. We found multiple promising combinations with synergistic anti-Ewing activity and experimentally validated the *in vitro* and *in vivo* efficacy of the combination of copanlisib, a PI3K inhibitor, and ribociclib, a CDK4/6 inhibitor for Ewing sarcoma. Our study supports the exploration of a PI3K inhibitor plus CDK4/6 inhibitor combination as a potential treatment strategy in Ewing sarcoma and provides a resource of additional combinations of drugs and targets for further validation in this disease.

## Materials and Methods

### Cell lines

All cell lines were provided by American Type Culture Collection (ATCC) or the Stegmaier laboratory at the Dana-Farber Cancer Institute. Lines were tested for mycoplasma contamination and verified by short tandem repeat profiling prior to drug screening. The cell lines were cultured at 37°C in a humid atmosphere that contained 5% CO2. A673 and EW8 cell lines were grown in DMEM (Corning catalog No 10-013- CV) supplemented with 10% FBS (Sigma-Aldrich), 1% sodium pyruvate (Gibco catalog No. 11360-070. TC32 and TC71 cell lines were grown in RPMI (Corning catalog No 10- 040-CV) supplemented with 10% FBS (Sigma-Aldrich), 1% sodium pyruvate (Gibco catalog No. 11360-070. All cells were cultured in the presence of Penicillin Streptomycin L-Glutamine 100X (Corning 30-009-CI).

### Single Agent Dose Response Curves

To identify the appropriate dose range of each single agent for the combination screen, we performed single-agent dose-response curves on A673 and TC32 Ewing cell lines across three classes of compounds: cell cycle inhibitors, TKI/mTOR inhibitors, and chemotherapeutic agents including. Copanlisib (catalog No. S2802), ribociclib (catalog No. S7440), prexasertib (catalog No. S7178), barasertib (catalog No. S1147), MK- 1775/adavosertib (catalog No. S1525), alisertib (catalog No. S1133), GSK-1070916A (catalog No. S2740), linsitinib (catalog No. S1091), dasatinib (catalog No. S1021), gemcitabine (catalog No. 1714), vincristine sulfate (catalog No. S1241), ulixertinib (catalog No. S7854), cabozantinib (catalog No. S1119), trametinib (catalog No. 2673), regorafenib (catalog No. S1178), LY-3023414 (catalog No. S8322), irinotecan (catalog No. S2217), NVP-AEW541 (catalog No. S1034), AZD-6738 (catalog No. S7693), torin-1 (catalog No. S2827), ZSTK-474 (catalog No. S1072), PF-562271 (catalog No. S2890), volasertib (catalog No. S2235), dinaciclib (catalog No. S2768) were all purchased from Selleck Chemicals. Palbociclib was purchased from Selleck Chemicals (catalog No. S4482) and Cell Signaling Technology (catalog No 47284S). SN-38 was purchased from Selleck Chemicals (catalog No. S4908) and Thermo Fisher Scientific (catalog No. NC0621047). Doxorubicin was purchased from Cell Signaling Technology (catalog No. 5927S), GSK-461364A (Sigma Aldrich SML1912).

45μL of cells suspended in media were transferred into 384-well cell culture plates (Corning 3570) at a density of 25,000 cells per mL of media using a BioTek Multiflo FX Multimode Dispenser. Then 5μL of media containing 10X of the target concentrations of the compounds of interest were robotically added to the cells by a Bravo Automated Liquid Handling Platform (Agilent) with a minimum of 4 replicates per concentration.

Compounds were added across a range of 10 concentrations and cells were also treated with the empty vehicle (most often DMSO) as a control. Initially, cells were treated for 48hrs with a range of concentrations across a 1:2 serial dilution starting at the highest concentration of 20μM. To measure cell viability after 48 hours of incubation, 10μL of Cell-TiterGlo (Promega, #G7572) was robotically dispensed to each well (BioTek, MultiFlo FX Multi-mode Dispenser, #7171013) and luminescence was measured using the FLUOstar Omega microplate reader (BMG Labtech) following a 10- minute incubation time. The half-maximal inhibitory concentration (IC50) for each compound was calculated using GraphPad Prism 5.0 (GraphPad Software, Inc). After initial results, we adjusted starting concentrations and the rate of serial dilution (for example, 3:4) for some compounds to identify an optimal range of concentrations which we defined as a range in which at least one concentration demonstrated almost no inhibitory effect on viability, at least one demonstrated 90% inhibitory concentration (IC90), and at least 3 concentrations impacted viability between these extremes and crossing the calculated IC50. This was not possible for all compounds. For example, single agent dasatinib had only a modest impact on cell viability in the TC32 cell line even at a concentration of 20μM.

### Small-molecule library drug combination screening

The combination matrix screening was performed by the National Center for Advancing Translational Sciences (NCATS) with the focused set of compounds in both A673 and TC32 cells. Plating of compounds in matrix format using acoustic droplet ejection and numerical characterization of synergy, additivity, and/or antagonism were conducted as described in our previous publication (26). The compounds were plated as a 10 × 10 dose-response combination matrix, and the concentration ranges were selected from single-agent dose-response curves generated from the qHTS. Compounds were acoustically dispensed (10nL/well) using an ATS-100 (EDC Biosystems) onto 1,536- well, white, solid-bottom, TC-treated plates. A673 or TC32 cells were subsequently added to the plates (1,000 cells/well in 5 µL) and incubated for 72 hours at 37 °C with 5% CO2 under 85% humidity. Cell viability was determined by the addition of 2.5 μL of CellTiter-Glo into each well. After 15 minutes of incubation at room temperature, each sample’s luminescence intensity was measured using a ViewLux reader. DMSO (20 nL) and bortezomib (20 nL at 2.3 mM) were used as negative and positive controls, respectively. Viability resulted from single-agent or combination was normalized to the controls. To identify combinations with synergistic anti-Ewing activity, multiple parameters of synergy were calculated including excess over highest single agent (HSA) as previously described (26).

### Validation of selected combinations

Selected synergistic combinations were then retested in five Ewing sarcoma cell lines (A673, TC32, EW8, TC71, and EWS502). 40μL of cells were seeded at a density of 17,000-25,000 cells per mL in 384-well white flat bottom tissue culture plates using a BioTek MultiFlo FX Multimode Dispenser. Each compound being tested was first prepared at a 10X concentration in media (across the same range as tested in the synergy screen) in 96-well plates with one compound titrated across columns (high to low concentrations) for “drug plate #1” and the other compound titrated by row for “drug plate #2”. Then 5μL of media containing 10X of the target concentration of the compound was robotically added to the cells by a Bravo Automated Liquid Handling Platform (Agilent). This was done for both drug plate #1 and #2 such that each concentration of compound #1 was tested against each concentration of compound #2 in 4 replicates.

After 48- and 72-hours of incubation, the Cell-TiterGlo luminescence assay was used as described above to quantify cell viability. Data from each individual well was converted to a fractional inhibition after first normalizing to the DMSO-treated replicates. To identify combinations with synergistic anti-Ewing activity, we then calculated the Combination index (CI) and Excess Over Bliss score for each agent as previously described (25,26). The Bliss independence model compares the observed effects of the drug combination to the expected effects, assuming the drugs act independently. CI values lower than 0.7 represent synergistic drug combinations, CI values between 0.7-1.3 indicate drugs that are additive, and CI values over 1.3 indicate antagonism.

### Protein analysis

#### Reverse-phase protein array analysis

EW8 cell lines were plated in 10 cm Petri dishes and treated the following day at approximately 30% confluency for 48 hours at the IC50 of copanlisib and/or ribociclib alone, which was determined in prior *in vitro* studies. One million cells from each treatment group were pelleted and submitted to the MD Anderson Cancer Center core facility for performance of the RPPA assay (https://www.mdanderson.org/research/research-resources/core-facilities/functional-proteomics-rppa-core.html) as described previously (27,28). Briefly, a panel containing 499 antibodies was used to measure protein abundance and phosphorylated protein levels from extracted protein lysates. Data were adjusted for set-to-set differences (batch effects) by normalizing identical control samples in this set with an invariant control sample set and applying the adjustment (differences in means × inverse of standard deviation ratio) to each corresponding data point. The resulting data were then transformed from log2 to linear values. Heatmaps were generated using the open-source software Morpheus (https://software.broadinstitute.org/morpheus). Values in the heatmap were mapped to colors using the minimum and maximum protein or phospho- protein levels of each row independently.

#### Western immunoblotting

Cells from A673, TC32, and EW8 cell lines were treated with the IC50 of each single agent, the combination, or vehicle control for 48 hours. One million cells were counted using the cell counter Countess II (Invitrogen) and pelleted for protein extraction with Cell Lysis Buffer (Cell Signaling Technology Catalog No. 9803S) supplemented with protease (Sigma Aldrich catalog No. 04906837001) and phosphatase inhibitor (Sigma Aldrich catalog No. 04906837001) tablets. The cell lysates were centrifuged at 16,000 x g for 10 minutes at 4°C following a 30-minute incubation on ice and the supernatant containing the protein was collected.

Protein concentrations were determined by the Bio-Rad Protein Assay Dye Reagent Concentrate protein assay according to the manufacturer’s instructions (Bio-Rad catalog No. 5000006). Protein samples were separated by standard SDS-PAGE method and transferred to PVDF membranes. Membranes were incubated with primary antibodies directed against AURKA (Cell Signaling Technology catalog No 3092S), AURKB (Cell Signaling Technology catalog No 3094S), phospho-S6 (S240/244) (Cell Signaling Technology catalog No 5364S), total S6 (Cell Signaling Technology catalog No 2217S), phospho-4EBP1 (T37/46) (Cell Signaling Technology catalog No 2855S), total 4EBP1 (Cell Signaling Technology catalog No 9644S), Cyclin B1 (Cell Signaling Technology catalog No 4138S), GAPDH (Cell Signaling Technology catalog No 5174S), vinculin (ABCAM catalog No AB129002). Horseradish peroxidase-conjugated secondary antibodies were used. Western blots were visualized by chemiluminescence using the Bio-Rad ChemiDoc MP instrument.

### RNA-sequencing

Ewing sarcoma cell lines A673, EW8, and TC32 were plated at a density of 250,000 cells per well in a 6 well plate and were treated with copanlisib, ribociclib, the combination or vehicle at the 48-hour IC50 for each agent. After trypsinizing, the treated cells were then harvested. The suspension was centrifuged at 800 x g in a refrigerated centrifuge for 5 minutes to decant culture media. The cell pellet was then washed with 10 mL cold PBS and spun down at 800 x g for 5 minutes at 4°C. Total RNA was extracted using the RNeasy Micro kit (Qiagen; Cat. No. 74004). For all samples, mRNA library preparation (poly A enrichment) and bulk RNA- sequencing on the Illumina platform (NovaSeq PE150) were performed at Novogene Bioinformatics Technology Co., Ltd. Samples were profiled in duplicate (A673) and triplicate (TC32 and EW8).

The sequencing reads were mapped to the GRCh37/hg19 human genome using STAR v2.7.2.b with standard parameters. Quality control tests for the aligned reads and for replicate consistency were performed using sample-sample clustering analysis, principal component analysis (PCA), and t-distributed Stochastic Neighbor Embedding (t-SNE) methods. Gene-level readings were summarized through the quantification of reads that aligned with the annotated reference of GRCh37/hg19 using the RSEM v1.3.1. Gene counts were normalized and utilized to quantify differential expressions between the experimental and control conditions using R package DESeq2 v1.40.2. The differential gene expression (DGE) values were also evaluated using the DESeq2 package, leveraging robust shrunken log_2_ (fold change) scores and approximate posterior estimation for the generalized linear model (GLM) coefficients (apeglm v 1.22.1) (29). The cutoffs for DGEs were | log_2_ (fold change)| ≥ 1 and adjusted P-value ≤ 0.05. Heatmaps representing the RSEM Log_2_(TPM+1) values (average of 3 replicates for each treatment arm) were generated using the open-source software Morpheus (https://software.broadinstitute.org/morpheus). Values in the heatmap are mapped to colors using the minimum and maximum of each row independently.

### Gene Set Enrichment Analysis (GSEA)

The GSEA v4.3.2 software was utilized to identify pathways and gene sets most enriched based on RNA sequencing data. Genes were ranked in the software based on Normalized Enrichment Score (NES), p-val, FDR q-val, and FWER p-val. The GSEA Preranked module was run across the collection of hallmark gene sets, curated gene sets, regulatory gene sets, and ontology gene set pathways available in the MSigDB v4.3.2 database with the significance cut-offs p-value 0.05 and FDR value 0.05. The MSigDB human collection pathways identified Hallmark gene sets and Reactome gene sets to be the most beneficial with 1692 gene sets analyzed by Reactome and 50 gene sets analyzed by Hallmark gene sets.

### Cell-cycle analysis and apoptosis assay

Cell death was measured using the Apoptosis Detection Kit-APC (eBioscience). After treatment, cells were trypsinized, collected and washed once with PBS and then resuspended with the provided binding buffer at 1×10^6^ cells/mL. The fluorochrome-conjugated Annexin V was first added to the cell suspension and incubated for 15 minutes at room temperature. After washing cells with 1X binding buffer, Propidium Iodide (PI) solution was added to the cell suspension. Fluorescence intensity was measured by flow cytometry within four hours (BD LSRFortessa Cell Analyzer) to differentiate apoptotic (Annexin V positive) from necrotic (PI positive and Annexin V negative) and live cells (negative for both). We excluded the necrotic population from our analysis.

For cell cycle analysis, cells were tryspinized, collected and washed with PBS. Cells were then fixed with 70% ethanol and incubated at -20C for at least two hours. After centrifugation at 800g for 5 minutes, the cell pellet was washed with PBS and spun again to remove the supernatant. Following an additional wash with the BD Pharmingen Stain Buffer (Catalog No. 554656), cells were incubated for 15 minutes with the BD Pharmingen PI/RNAse staining buffer (catalog No. 550825) at 2×10^6^ cells per 1mL buffer. Incubation was performed at room temperature and protected from light. No washing step was used before flow cytometry analysis.

A minimum of 50,000 stained cells were analyzed for all flow cytometry experiments. Flow data analysis was performed using the Omiq software (https://app.omiq.ai/login). All statistical analyses were performed using GraphPad Prism 5.0 software (GraphPad Software Inc., La Jolla, CA, USA). We used the One-way ANOVA nonparametric multiple comparisons test to compare the average percentage of apoptotic cells between each treatment group.

### Ewing sarcoma tumor formation and *in vivo* drug combination studies

#### Tolerability study

All animal studies were performed according to the Dana-Farber Cancer Institute Institutional Animal Care and Use Committee (IACUC)- approved protocol 17-028. NOD-SCID NSG mice were obtained from the Jackson Laboratory (USA). To confirm the tolerability of the drug combination, 5 male mice were treated for 5 days with the combination of ribociclib (oral gavage, 75mg/kg in 0.5% methylcellulose, CAS# 1211441-98-3) and copanlisib (intraperitoneal injection, 14mg/kg in 5% trifluoracetic acid and 0.9% NaCl, CAS# 1032568-63-0). Body weights were measured daily during dosing period, and all mice were monitored for an additional 7 days after dosing ended.

#### Efficacy study

To evaluate the efficacy of the combination of copanlisib and ribociclib, mice were injected with five million TC32 cells subcutaneously into the flanks of 8-weeks old female mice. When subcutaneous tumor volume reached 100-200 mm^3^, mice were randomized into four treatment groups: vehicle; ribociclib alone at 75mg/kg/day daily; copanlisib alone at 14mg/kg every other day; and the combination. Each group was treated for 15 consecutive days (Efficacy cohort, N=40 mice, ten mice per treatment arm). Tumors were measured by caliper every three days throughout treatment until the mice either reached their pre-defined endpoint of morbidity or tumors reached 2000- 2500 mm^3^.

#### Survival study

To determine the impact of the novel combination on survival, mice were injected with TC32 cells as described above and monitored by caliper measurements until tumor volumes reached 100-200 mm^3^ at which point mice were randomized into the four treatment groups described above (ten mice per arm). Mice were then treated according to assigned arm for 42 consecutive days. Mice were euthanized whenever the animal reached study endpoints which included signs of morbidity, a tumor size of approximately 2000-2500 mm^3^, or evidence of trauma due to repeated oral gavage trauma (confirmed by necropsy). For mice sacrificed during ongoing therapy due to gavage trauma, tumor tissue was collected and snap frozen.

#### Statistical analysis

All statistical analyses were performed using GraphPad Prism 5.0 software (GraphPad Software Inc., La Jolla, CA, USA). We used the Mann-Whitney nonparametric test to compare the average tumor size in each treatment group. To assess differences in survival in each mouse study, we performed Kaplan-Meier survival analysis and the Mantel-Cox test to compare the survival distributions between the four treatment groups.

#### Extraction of proteins from mouse tissues

Protein was extracted from tumors harvested from mice that experienced gavage trauma during ongoing therapy. 100μL of cold lysis buffer was added to tumor tissue which was then triturated 10-20 times using 1.5mL pestles. The mixture was incubated on ice for 30 minutes and then centrifuged at 16,000 x g for 10 minutes at 4C. Protein expression was evaluated by Western immunoblotting as described above. Membranes were incubated with primary antibodies directed against AURKA, AURKB, phospho-S6 (S240/244), total S6, Ki67 (Cell Signaling Technology, catalog No 16667), GAPDH, vinculin, phospho-4EBP1 (T37/46), total 4EBP1. Horseradish peroxidase-conjugated secondary antibodies were used. Western blots were visualized by chemiluminescence using the Bio-Rad ChemiDoc MP instrument.

## Results

### High-throughput small-molecule screening identifies PI3K and CDK4/6 inhibitors as highly synergistic in Ewing Sarcoma

Prior work in Ewing sarcoma suggests that the combination of inhibitors that downregulate mTOR pathway activity with agents that perturb cell-cycle regulation have synergistic anti-Ewing activity (15–24). To identify new candidate combinations of agents that might be readily translatable to clinical trials, we set out to conduct an *in vitro* combination screen in Ewing sarcoma cell lines of anti-cancer agents from these two drug classes. Through a literature review, advice from clinical trial experts on our team, and by searching clinicaltrials.gov, we identified compounds from these drug classes that were available in the NCATS extensive compound library and were either already FDA-approved or in late-phase clinical trials. We chose 12 kinase inhibitors known to impact mTOR pathway activity, 10 compounds that inhibit cell-cycle regulators, 5 chemotherapy agents, and one ATR inhibitor for our study. (**Figure 1A and Supplemental Table 1**).

**Figure 1:**
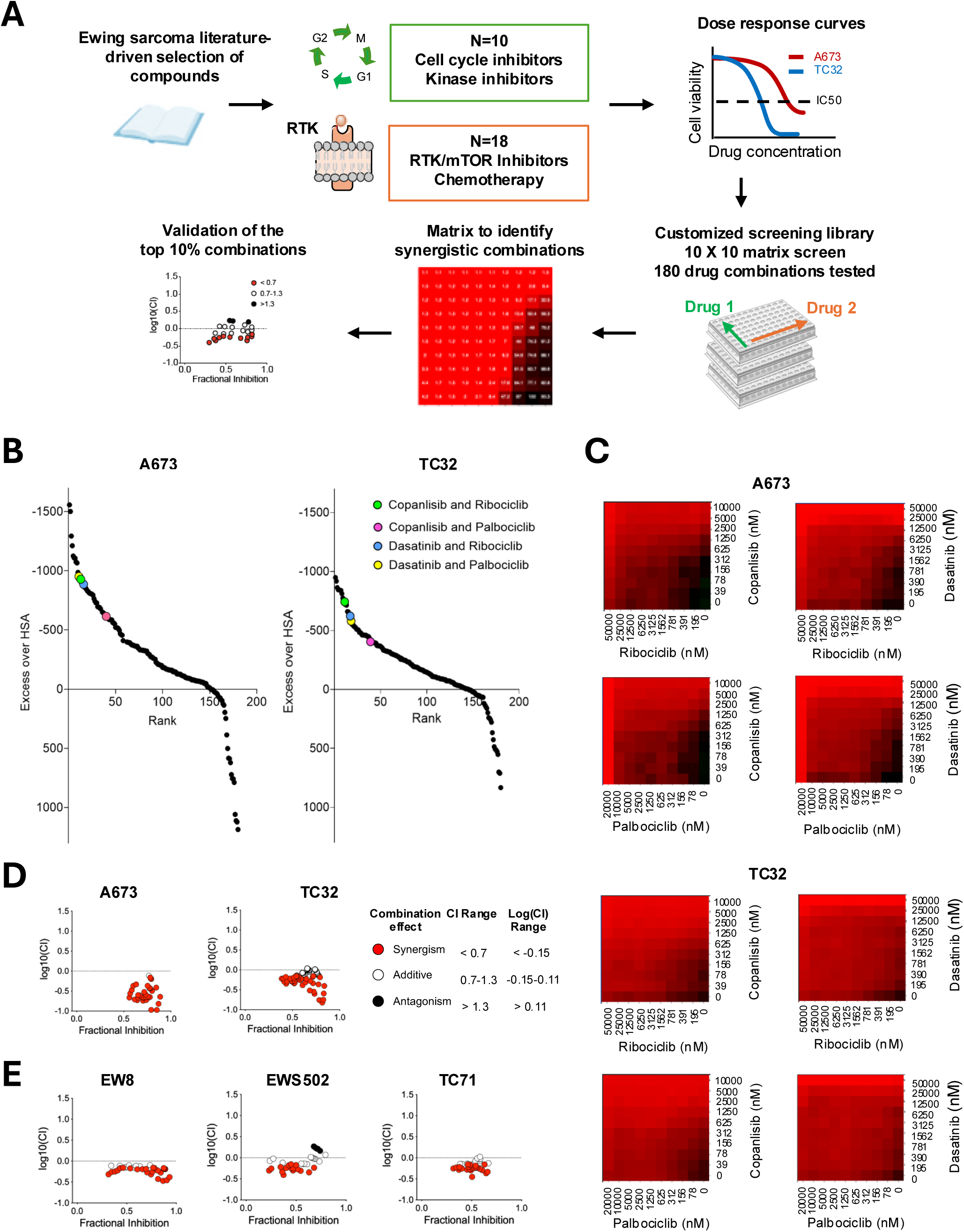
Custom small-molecule screening identifies PI3K and CDK4/6 inhibitors as highly synergistic in Ewing Sarcoma cells. **A-** Schema of custom drug screen to identify synergistic anti-Ewing drug combinations. Literature review and expert opinion solicited to select 28 FDA-approved and late-phase agents targeting PI3K/AKT/mTOR pathway, cell cycle regulation, and chemotherapy agents. Single-agent dose response curves conducted in Ewing lines for each compound to identify optimal screening doses. 180 combinations were then screened using a 10×10 matrix of disease for each combination. Synergistic combinations identified from the screen were then validated in replicate viability experiments using a similar matrix design. **B-** Scatter plots depicting the results of synergy screen in Ewing cell lines. Excess over HAS values are plotted for each line on the y-axis with rank of these values plotted on x- axis. Negative values indicate synergy and positive values indicate antagonism. **C-** Heatmaps of 10×10 matrices of select synergistic combinations from screening data. Red indicates the greatest inhibition of cell viability and black indicates no inhibition. **D-E** Scatter plots of the log10 normalized combination index (CI) values versus the fractional inhibition of cell viability from validation experiments in Ewing sarcoma cell lines.

To optimize the performance of the screen, we first performed single-agent dose response curves for each of the 28 selected drugs using the A673 and TC32 Ewing cell lines **(Figure 1A and Supplemental Table 1)**. Dose ranges from these experiments were then utilized to develop a customized screening library to identify combinations with synergistic anti-Ewing activity. In this screen, the NCATS team arrayed each compound across a range of 10 doses for testing in Ewing cell lines. Each kinase inhibitor known to regulate the mTOR pathway was screened against each cell-cycle inhibitor, chemotherapy, and the ATR inhibitor resulting in 180 combinations. Each of these combinations was screened for synergy in both the A673 and TC32 cell lines **(Figure 1A)**. Cells treated with each combination were treated for 72 hours and viability was measured. To identify synergistic combinations, viability measurements were used to calculate the excess over HSA model ranked to identify combinations with the most doses demonstrating synergistic inhibition of Ewing sarcoma cell viability (**Figure 1B and Supplemental Figure 1**). Consistent with prior literature, combinations of IGF-1R inhibitors with agents targeting CDK4/6 and other cell cycle regulators, scored in the top 10% of combinations **(Supplemental Table 2 and 3)** (20). AURKA/B inhibitors in combination with multiple agents that downregulate mTOR pathway activity (IGF-1R, PI3K, and pan-tyrosine kinase inhibitors; pan-TKI) also scored as synergistic combinations consistent with prior studies **(Supplemental Table 2 and 3)** (25). Of the remaining combinations that scored in the top 10%, only combinations of copanlisib, a PI3K inhibitor, or disatinib, a pan-TKI, with CDK4/6 inhibitors were composed of FDA-approved drugs that could be most easily adapted to a clinical trial **(Figure 1C and Supplemental Figure 2)**. We validated these combinations in the A673 and TC32 cell lines and found that copanlisib with ribociclib demonstrated the most robust synergy across a range of drug concentrations **(Figure 1D and Supplemental Figure 3)**. We then confirmed that copanlisb and ribociclib demonstrated *in vitro* synergy in additional Ewing sarcoma cell lines **(Figure 1E)**.

### Combination PI3K and CDK4/6 inhibition induces cell cycle dysregulation and apoptosis

We next profiled the impact of PI3K and CDK4/6 inhibition on protein expression and kinase signaling on Ewing sarcoma cells using an established reverse-phase protein array (RPPA) assay (27,28). EW8 Ewing sarcoma cells treated for 48 hours with either DMSO, ribociclib alone, copanlisib alone, or both drugs together at concentrations corresponding to the IC_50_ (ribociclib = 15 nmol/L and copanlisib = 0.5 nmol/L), were subjected to protein array analysis. Treatment of EW8 cells with copanlisib and the combination demonstrated loss of phosphorylation of downstream effectors of the PI3K/AKT pathway such as the proteins S6 and 4E-BP1, despite upregulation of phosphorylation of the mTOR protein **(Figure 2A)**. We also observed downregulation of proteins involved in cell cycle regulation after treatment with the combination, including Aurora kinases A and B and the CDK1 (Cyclin-dependent kinase 1)-Cyclin B1 complex **(Figure 2B)**. Downregulation of cell cycle-associated proteins was lowest in EW8 cells treated with the combination.

**Figure 2:**
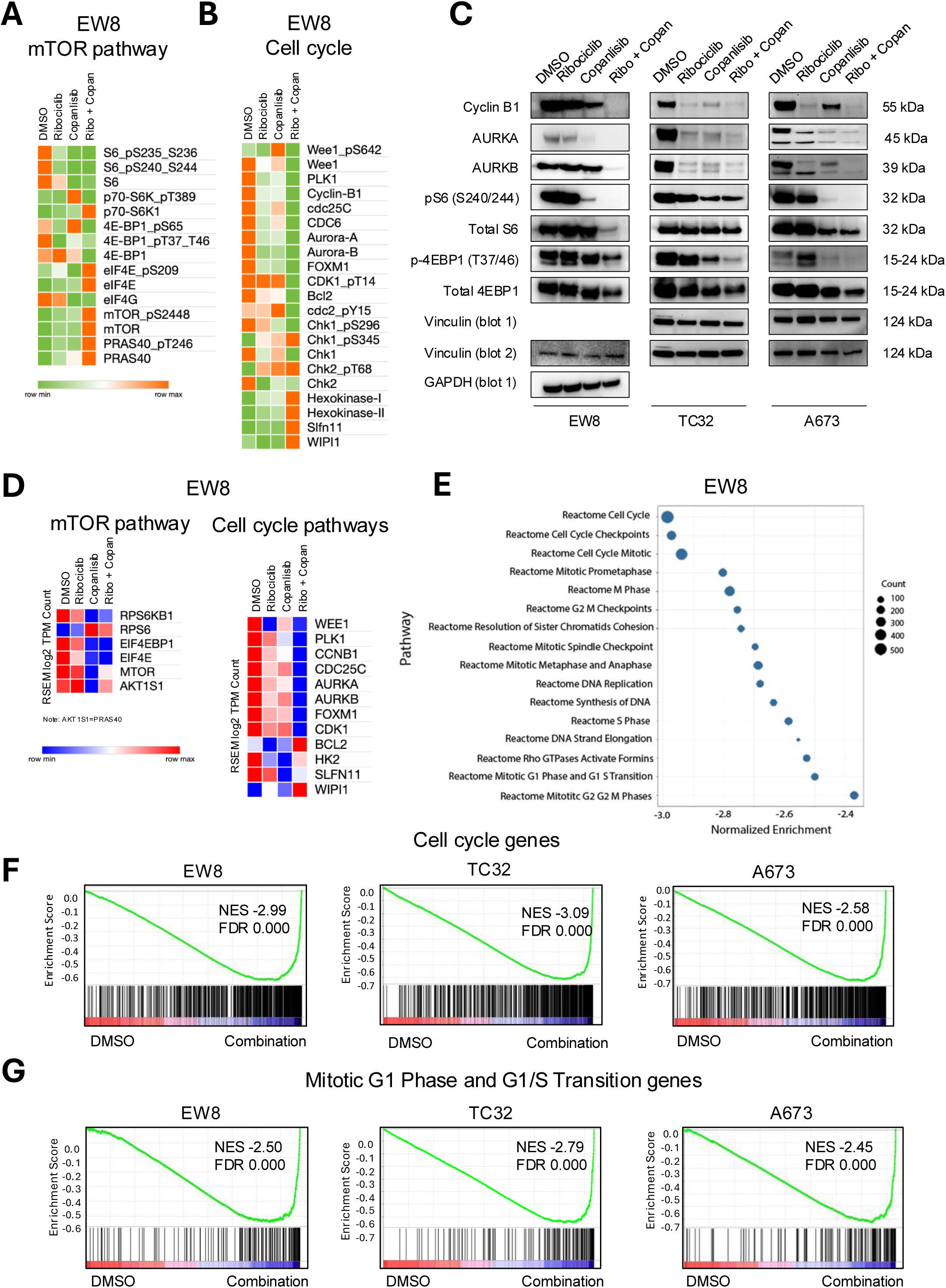
Gene expression changes in Ewing sarcoma cells treated with PI3K and CDK4/6 inhibitors. **A-B-** Heatmap of normalized expression levels of (A) mTOR pathway and (B) cell cycle proteins from RPPA analysis in EW8 Ewing sarcoma cells after 48-hours incubation with the indicated compounds. **C-** Representative western blots of cell cycle-related signaling pathways in Ewing cell lines after 48 hours of treatment with ribociclib (IC50), copanlisib (IC50) and the combination. Vinculin and GAPDH were used as loading controls. **D-** Heatmap visualization gene expression (transcript per million) data from RNA-Seq of EW8 Ewing cells treated for 48-hours with the indicated compounds. Genes listed on the heatmap were selected based on RPPA results. Each column shows average data from three independent replicates. **E-** Dot plot of gene signatures from the Reactome Set of gene signatures (MSigDB) that are significantly enriched for gene expression changes observed in EW8 cells treated with the combination versus DMSO. The signatures are sorted based on the normalized enrichment score on the x-axis. The size of the dot represents the number of genes in the signature. All enrichments had a P-value <0.0001. **F-G** GSEA plots of running enrichment scores for the Reactome set (F) Cell Cycle gene signature and (G) Mitotic G1 Phase and G1/S Transition gene signature significantly enriched in Ewing cell lines treated with the combination (vs. DMSO).

We validated these findings using western immunoblotting across three Ewing cell lines **(Figure 2C)**. We showed a stronger downregulation of phospho-S6 (S240/244), phospho-4EBP1 (T37/46), and Cyclin B1 in cells treated with the combination compared to cells treated with single agents or DMSO. The expression of Aurora Kinase A or B (AURKA/B), necessary for normal proliferation in Ewing sarcoma cells, was also strongly reduced in cells treated with the combination.

To further validate findings from the RPPA assay, we performed RNA-seq profiling across three Ewing cell lines. We observed a significant downregulation of genes involved in mTOR-regulated proliferation and cell cycle regulation in cells treated with the combination compared to DMSO or cells treated with single agents, confirming our proteomic results **(Figure 2D and Supplemental Figure 4)**. Gene Set Enrichment Analysis (GSEA) of this RNA-Seq data using the Molecular Signature Database (MSigDB) demonstrated that the combination therapy triggered a significant enrichment for signatures of mTORC1 signaling, E2F targets, cell cycle regulation, G2M checkpoint, mitotic spindle, and mitosis **(Figure 2E-G and Supplemental Figure 5)**.

To evaluate the phenotypic effects of treatment of Ewing cell lines with PI3K and CDK4/6 inhibition, we performed apoptosis and cell cycle analysis using a flow cytometry-based workflow **(Figure 3A)**. We treated TC32, EW8, TC71 and A673 cell lines with either control DMSO, ribociclib alone, copanlisib alone or combination of the two at IC_50_ (ribociclib = 15 nmol/L and copanlisib = 0.5 nmol/L) for 48 hours. We found that the CDK4/6 inhibitor, ribociclib, induced a G0/G1 arrest in the majority of cells in all Ewing lines while the PI3K inhibitor, copanlisib, had a more modest impact on cell cycle arrest **(Figure 3B-E)**. We also showed that ribociclib had minimal impact on apoptosis while treatment with copanlisib in combination with ribociclib had the most significant increase in apoptosis compared to DMSO **(Figure 3B-E)**.

**Figure 3:**
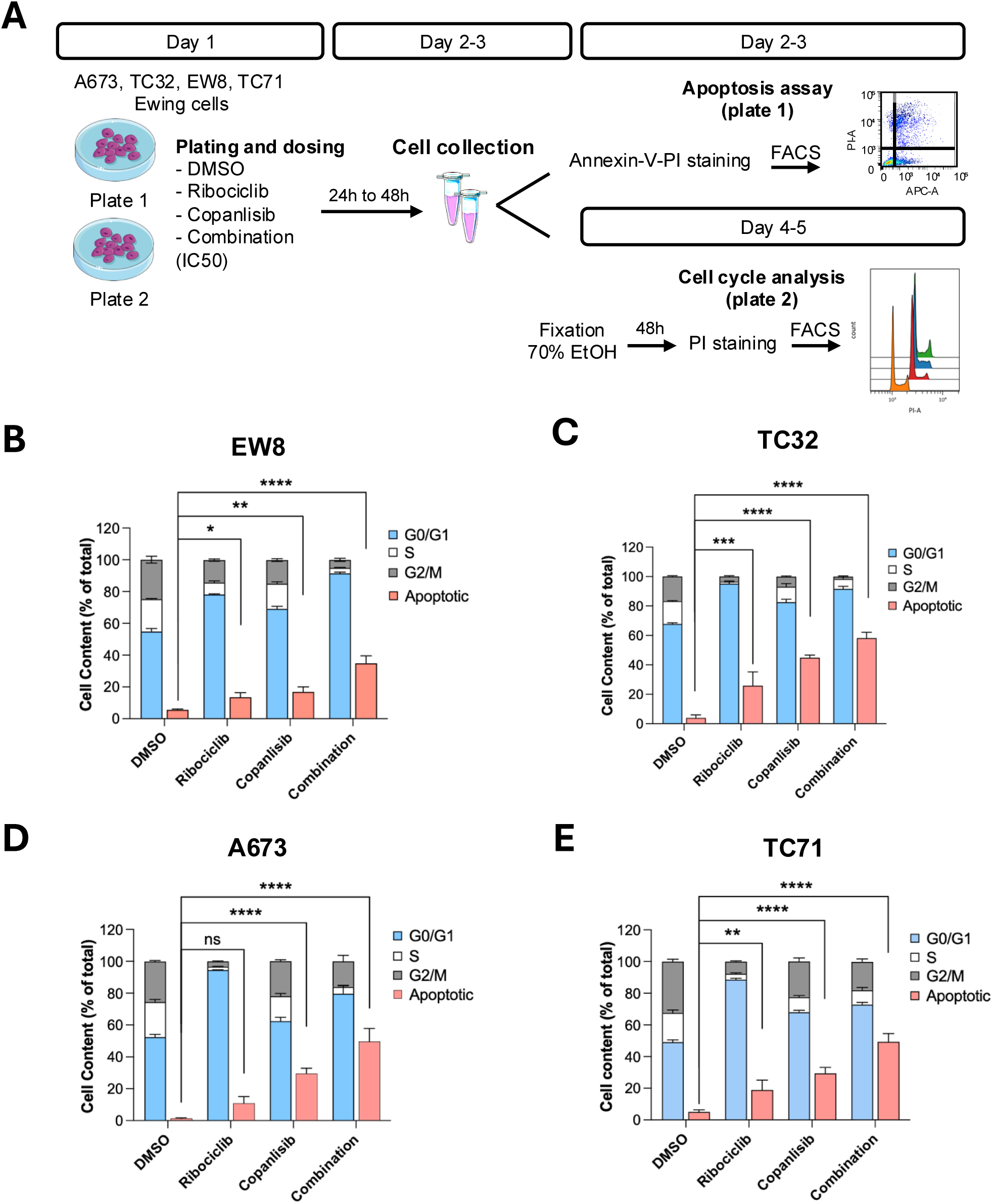
*In vitro* functional validation of the novel combination of PI3K inhibitors with CDK4/6 inhibitors in Ewing cell lines. **A-** Schematic displaying the approach to phenotypic profiling of Ewing cell lines treated with ribociclib, copanlisib, and the combination vs. vehicle control. **B-E** Cell cycle and apoptosis analysis following 48-hours of treatment with DMSO or the indicated compounds the IC50 of the single agents or the combination of both in Ewing cell lines. Bars show the mean of 4 technical replicates with error bars representing the standard error of the mean (SEM). Biological replicates were performed to confirm the results. **** p-value <0.0001.

### Combined PI3K and CDK4/6 inhibition significantly impairs Ewing sarcoma tumor progression and improves survival for *in vivo* models

To assess the *in vivo* tolerability associated with the combination of ribociclib and copanlisib in NSG immunocompromised mice, animals were treated with ribociclib by daily via oral gavage at 75mg/kg/day, a dose previously reported in the literature in Ewing sarcoma (20) and intraperitoneal injection of copanlisib was given every other day at the maximum tolerated dose in mice (14mg/kg). No significant weight loss was observed after 5 days of treatment indicating that the combination therapy was well tolerated **(Figure 4A),** confirming published data in breast cancer (30). Notably, the short-term nature of this study was not designed to capture the toxicity of prolonged daily gavage that was later experienced in our survival study.

**Figure 4:**
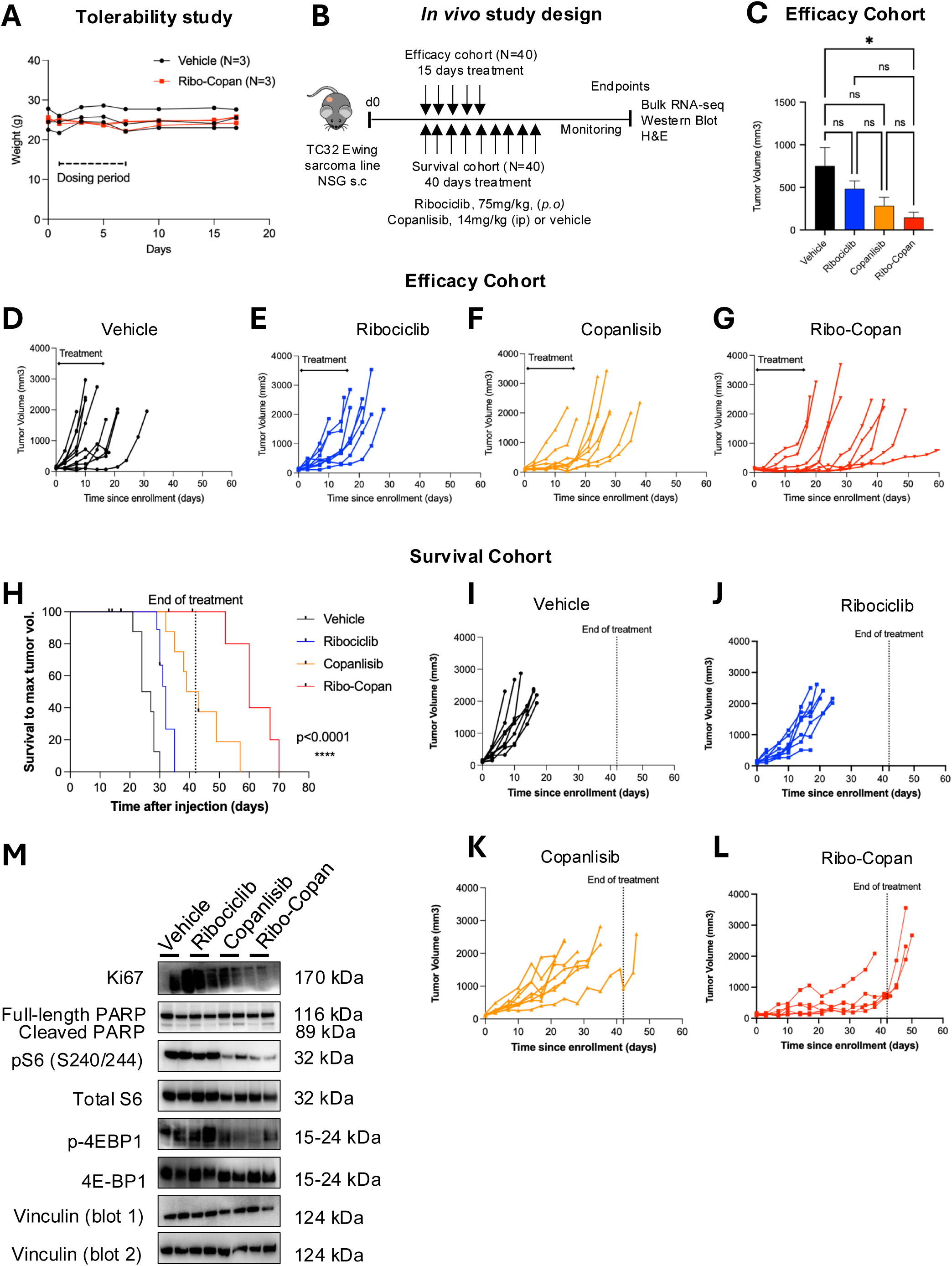
Combined PI3K and CDK4/6 inhibition significantly prolonged survival in Ewing sarcoma mouse models. **A-** Plot of individual mouse weights from the vehicle- (n=3) and combination- treated (n=3) animals during the tolerability study. Black dotted line showing the treatment period. **B-** Schematic of enrollment and treatment strategy for efficacy study (panels C-G) and survival study (panels H-L) after subcutaneous transplantation of TC32 cells in SCID immunocompromised mice. Treatment was initiated when tumor volume reached 100mm^3^. **C-** Bars represent mean caliper measurements of tumor volume in mice treated with vehicle, single agents, and the combination in the efficacy cohort. Error bars indicate SEM. Treatment arms compared by Mann-Whitney nonparametric test. ns non- significant; * p-value < 0.05. **D-G** Spider plots showing tumor size by caliper measurements in vehicle, single agent and combination treated mice from the efficacy cohort (n=10 mice per arm). Solid black line showing the treatment period. **H-** Kaplan-Meier survival curve for mice treated for 42 days with the indicated therapy after establishment of TC32 xenograft tumors. Significance determined by log-ranked test. **I-L-** Spider plots showing tumor size by caliper measurements in vehicle, single agent and combination treated mice from the survival cohort (n=10 mice per arm). Dotted vertical black line indicates the end of the treatment period. **M-** Western blot analysis of tumor lysates extracted from mice treated in the survival study with vehicle, single agent, or in combination treatment and sacrificed during the treatment period. Vinculin was used as loading control.

To determine the impact of the combination therapy on short-term tumor proliferation, we next implanted TC32 cells subcutaneously in NSG mice (n = 40), allowed tumors to reached 100-200 mm3 and then treated daily for 15 consecutive days **(Figure 4B Supplemental Figure 6A)**. Tumors in mice treated with vehicle progressed rapidly and several were euthanized prior to completing 15 days of vehicle **(Supplemental Table 4)**. Tumor sizes at day 7 were significantly smaller in the combination arm than in the vehicle arm **(Figure 4C-G and Supplemental Table 5).** While this experiment was not designed to demonstrate a survival benefit, the mice treated with the combination had significantly longer survival than vehicle-treated mice, while single-agent therapy had no significant impact on survival **(Supplemental Figure 6B)**.

To measure the impact of this treatment combination on survival, we again treated TC32-engrafted mice with vehicle, single-agent, or combination therapy. In this experiment mice were treated for up to 42 days. Six mice, across treatment arms, were euthanized for gavage-related trauma or weight loss greater than 10% of their original weight **(Supplemental Figure 6C and Supplemental Table 6)** and were considered censored at the time of sacrifice in the survival analysis. Mice in the combination arm had delayed tumor progression and significantly improved survival over mice treated with vehicle or single-agent therapy **(Figure 4H-L)**. In fact, only one mouse in the combination arm demonstrated tumor progression during therapy while all mice, with the exception of one mouse from the copanlisib arm, reached the maximum tumor size before treatment was completed. All tumors progressed after the combination treatment ended.

Finally, we collected mouse tumor tissues during the treatment period for downstream signaling analysis from mice sacrificed due to gavage trauma/toxicity. Western blot analysis revealed a stronger decrease of phospho-S6 (S240/244), phospho-4EBP1, and the proliferative index Ki67 in tumors treated with copanlisib alone and the combination, compared to vehicle- or ribociclib-treated tumors **(Figure 4M)**. We observed unchanged level of PARP expression in tumor lysates across all treatment groups. Overall, our *in vivo* studies highlighted the benefit of the combination therapy to inhibit Ewing tumor growth and prolong survival.

## Discussion

The fusion protein EWSR1::FLI1 is recognized as the primary oncogenic driver in Ewing sarcoma. Modulating transcription factors pharmacologically has been historically difficult (14,31), and effective inhibitors specifically targeting EWSR1::FLI1 have not yet been clinically successful. One putative inhibitor of EWSR1::FLI1 activity, TK216, was recently shown to also inhibit tubulin, potentially accounting for its anti-Ewing activity, although direct inhibition of the chimeric protein may also contribute (32–35). Other attempts to perturb the activity of EWSR1::FLI1 oncoprotein in Ewing sarcoma have focused on inhibition of epigenetic regulatory molecules, such as the lysine-specific demethylase 1 (LSD1) gene (36,37). While *in vitro* and xenograft studies demonstrated that small molecule inhibition of LSD1 modulates the transcriptional activity of EWSR1::FLI1 proteins in Ewing sarcoma (37), a phase II study of seclidemstat failed to demonstrate significant efficacy in Ewing sarcoma (38). Trabectedin was recently reported to have an early efficacy signal in a phase II trial in Ewing sarcoma (39) and is thought to induce the sequestration of EWS/ETS chimeric proteins away from chromatin (40–42). This agent will require further testing in clinical trials and will likely require combination with other agents to elicit cures.

Beyond the fusion, we and others have shown that these tumors harbor few recurrent genomic aberrations and no druggable mutated kinases (6,11,12). This underscores the need for additional alternative therapeutic approaches to effectively combat this aggressive pediatric cancer. Numerous studies have reported that Ewing sarcoma proliferation is dependent on the activity of the mTOR pathway and several studies suggest that combining inhibition of the mTOR pathway with perturbation of cell-cycle regulation may have potent anti-Ewing activity (16,20–22,25). Despite the promising results of several early-phase single-agent trials (43), recent studies of IGF-1R inhibition in combination with chemotherapy or cell-cycle inhibitors failed to demonstrate compelling efficacy (4,44). Previous work from our group also demonstrates that inhibition of focal adhesion kinase (FAK) downregulates mTOR activity and has synergistic anti-Ewing activity when combined with Aurora kinase inhibition (21,25). Clinical trials testing these agents for patients with Ewing sarcoma have not been feasible due to a lack of access to agents. In fact, neither FAK nor AURK inhibitors have yet to reach FDA-approval for any indication.

Several promising initiatives such as the Pediatric Dependency Map (45) have been supported by the Moonshot Pediatric Cancer Challenge to identify new targetable dependencies in Ewing sarcoma and new direct inhibitors of the EWSR1::FLI1 oncoprotein. However, patients who have experienced relapse after initial diagnosis remain in urgent need of impactful therapeutic options and these new discovery efforts may take many years to result in clinical trials of novel agents. The track record of rapid accrual to early phase trials in Ewing sarcoma highlights the need for new options for these patients. In this study, we endeavored to identify novel combinations of targeted therapies using a high-throughput *in vitro* drug screen enriched for FDA-approved drugs and late-phase anti-cancer agents. We identified multiple candidate combinations with synergistic anti-Ewing activity and prioritized the validation of copanlisib and ribociclib due to their status as FDA-approved drugs and known individual tolerability in pediatric trials (NCT03458728, NCT01747876) **(Supplemental Table 7)**.

We demonstrated that the combination of copanlisib and ribociclib had synergistic anti-Ewing activity in a panel of five Ewing sarcoma cell lines, impaired tumor proliferation in a mouse xenograft model, and significantly improved survival of these mice compared to single-agent therapy when given over a six-week period. Due to the trauma of daily oral gavage, we were unable to test the effects of continual treatment with this combination beyond this two-month window, but tumor progression was largely abrogated until combination therapy was discontinued.

We also observed that single agent ribociclib had little effect on viability *in vitro* or on tumor proliferation in mice but did induce a striking cell cycle arrest consistent with the known mechanism of action of this agent and its characterization as a cytostatic agent in the clinic. However, treatment with copanlisib induced apoptosis *in vitro* and reduced tumor proliferation *in vivo*. This cytotoxic effect was significantly enhanced when given in combination with ribociclib. Transcriptomic and proteomic profiling of treated Ewing sarcoma cells demonstrated that this combination enhanced suppression of the activity of mTOR pathway effector molecules, S6 and 4EBP, over single-agent therapy, while also perturbing cell-cycle regulation.

While several studies have shown the potential efficacy of combining mTOR inhibition with perturbation of cell-cycle regulation in Ewing sarcoma, this study identifies and validates the efficacy of two FDA-approved, commercially accessible, and clinically tolerable drugs that could be immediately applied to a clinical trial for patients with this disease. Furthermore, our drug screen has highlighted multiple additional candidate combinations of agents, including multiple TKIs with CDK4/6 and WEE1 inhibitors, as additional agents that could be validated for therapeutic trials. We believe this screening approach could also be expanded to other rare treatment-refractory pediatric cancers for which there is a striking shortage of experimental therapies available for patients with relapsed disease.

## Acknowledgments

Rally Foundation Research Grant, Collaborative Pediatric Cancer Research Awards Program, Pediatric Cancer Research Foundation, Nathaniel Cavallo Fund, and Team Fernando’s Fight (BDC). Parts of this work were also supported by the Intramural Program of the National Center for Advancing Translational Sciences, National Institutes of Health (MH).

Illustrations were created using images from Servier Medical Art, licensed under a Creative Commons BY 4.0 (https://smart.servier.com).

## Author contributions

AFB, MH, SGD, and BDC conceived the study and supervised research. MAICDLS, HFG, COH, AAQ, BDC, SM, KK, and AFB designed and performed experiments. MAICDLS, HFG, AAQ, AFB, MT, and BDC analyzed the data. YQZ, MS, and MH performed the drug screen. All authors wrote, reviewed, edited and approved the manuscript.

## Declaration of interests

SGD reports consulting fees from Amgen, Bayer, InhibRx, and Jazz and travel expenses from Loxo, Genentech, and Salarius. DSS reports consulting: Boehringer Ingelheim.

## Supplemental Figure Legend

**Supplemental Figure 1: Drug matrix screen of 180 combinations.** Heatmaps of excess over HSA data of 180 combinations used to treat A673 and TC32 cells to identify synergistic combinations with anti-Ewing activity. The most synergistic (most negative) values are plotted as blue and antagonistic (more positive) values are plotted in yellow.

**Supplemental Figure 2: Additional drug screening results with compounds from the top 10 percent of the 180 combinations tested.**

Heatmaps of 10×10 matrices of select synergistic combinations from screening data from (A) A673 and (B) TC32 treated cells. Red indicates the greatest inhibition of cell viability and black indicates no inhibition.

**Supplemental Figure 3: Combination index results with CDK4/6 inhibitors in combination with pan-kinase or PI3K inhibitors.**

Scatter plots of the log10 normalized combination index (CI) values versus the fractional inhibition of cell viability from validation experiments in Ewing cell lines treated with the indicated combinations.

**Supplemental Figure 4: Heatmap visualization of gene expression changes in Ewing cells treated with CDK4/6 and PI3K inhibitors.** Heatmap visualization gene expression (transcript per million) data from RNA-Seq of (A) TC32 and (B) A673 Ewing cells treated for 48-hours with the indicated compounds. Genes listed on the heatmap were selected based on RPPA results. Each column shows average data from three independent replicates.

**Supplemental Figure 5: Gene set enrichment analysis of ribociclib and copanlisib combination in Ewing sarcoma cells**

**A-** GSEA plots of running enrichment scores for the the indicated gene signatures significantly enriched in Ewing cell lines treated with the combination (vs. DMSO).

**B-** Dot plot of gene signatures from the Reactome Set of gene signatures (MSigDB) that are significantly enriched for gene expression changes observed in the indicated Ewing cell lines treated with the combination versus DMSO. The signatures are sorted based on the normalized enrichment score on the x-axis. The size of the dot represents the number of genes in the signature. All enrichments had a P-value <0.0001.

**Supplemental Figure 6: Weight monitoring and survival analysis of mice enrolled in the efficacy and survival studies.**

**A-** Plot of individual mouse weights from the animals treated on in the efficacy cohort. Black dotted line shows the end of the treatment period.

**B-** Kaplan-Meier survival curve for mice treated in the efficacy cohort for 15 days with the indicated therapy after establishment of TC32 xenograft tumors. Significance determined by log-ranked test.

**C-** Plot of individual mouse weights from the animals treated on in the survival cohort. Black dotted line shows the end of the treatment period.

## Bibliography

1. Leavey PJ, Laack NN, Krailo MD, Buxton A, Randall RL, DuBois SG, et al. Phase III Trial Adding Vincristine- Topotecan-Cyclophosphamide to the Initial Treatment of Patients With Nonmetastatic Ewing Sarcoma: A Children’s Oncology Group Report. J Clin Oncol Off J Am Soc Clin Oncol. 2021;39:4029–38.

2. Leavey PJ, Mascarenhas L, Marina N, Chen Z, Krailo M, Miser J, et al. Prognostic factors for patients with Ewing sarcoma (EWS) at first recurrence following multi-modality therapy: A report from the Children’s Oncology Group. Pediatr Blood Cancer. 2008;51:334–8.

3. Stahl M, Ranft A, Paulussen M, Bölling T, Vieth V, Bielack S, et al. Risk of recurrence and survival after relapse in patients with Ewing sarcoma. Pediatr Blood Cancer. 2011;57:549–53.

4. DuBois SG, Krailo MD, Glade-Bender J, Buxton A, Laack N, Randall RL, et al. Randomized Phase III Trial of Ganitumab With Interval-Compressed Chemotherapy for Patients With Newly Diagnosed Metastatic Ewing Sarcoma: A Report From the Children’s Oncology Group. J Clin Oncol Off J Am Soc Clin Oncol. 2023;41:2098–107.

5. Marina NM, Liu Q, Donaldson SS, Sklar CA, Armstrong GT, Oeffinger KC, et al. Longitudinal follow-up of adult survivors of Ewing sarcoma: A report from the Childhood Cancer Survivor Study. Cancer. 2017;123:2551–60.

6. Crompton BD, Stewart C, Taylor-Weiner A, Alexe G, Kurek KC, Calicchio ML, et al. The Genomic Landscape of Pediatric Ewing Sarcoma. Cancer Discov. 2014;4:1326–41.

7. Campbell BB, Light N, Fabrizio D, Zatzman M, Fuligni F, de Borja R, et al. Comprehensive Analysis of Hypermutation in Human Cancer. Cell. 2017;171:1042–1056.e10.

8. Anderson ND, de Borja R, Young MD, Fuligni F, Rosic A, Roberts ND, et al. Rearrangement bursts generate canonical gene fusions in bone and soft tissue tumors. Science. 2018;361:eaam8419.

9. Lawrence MS, Stojanov P, Polak P, Kryukov GV, Cibulskis K, Sivachenko A, et al. Mutational heterogeneity in cancer and the search for new cancer genes. Nature. 2013;499:214–8.

10. Gröbner SN, Worst BC, Weischenfeldt J, Buchhalter I, Kleinheinz K, Rudneva VA, et al. The landscape of genomic alterations across childhood cancers. Nature. 2018;555:321–7.

11. Tirode F, Surdez D, Ma X, Parker M, Le Deley MC, Bahrami A, et al. Genomic landscape of Ewing sarcoma defines an aggressive subtype with co-association of STAG2 and TP53 mutations. Cancer Discov. 2014;4:1342–53.

12. Brohl AS, Solomon DA, Chang W, Wang J, Song Y, Sindiri S, et al. The Genomic Landscape of the Ewing Sarcoma Family of Tumors Reveals Recurrent STAG2 Mutation. PLoS Genet. 2014;10:e1004475.

13. Ma X, Liu Y, Liu Y, Alexandrov LB, Edmonson MN, Gawad C, et al. Pan-cancer genome and transcriptome analyses of 1,699 pediatric leukemias and solid tumors. Nature. 2018;555:371–6.

14. Dang CV, Reddy EP, Shokat KM, Soucek L. Drugging the “undruggable” cancer targets. Nat Rev Cancer. 2017;17:502–8.

15. Palombo R, Passacantilli I, Terracciano F, Capone A, Matteocci A, Tournier S, et al. Inhibition of the PI3K/AKT/mTOR signaling promotes an M1 macrophage switch by repressing the ATF3-CXCL8 axis in Ewing sarcoma. Cancer Lett. 2023;555:216042.

16. Truong DD, Lamhamedi-Cherradi S-E, Ludwig JA. Targeting the IGF/PI3K/mTOR pathway and AXL/YAP1/TAZ pathways in primary bone cancer. J Bone Oncol. 2022;33:100419.

17. Seligson ND, Maradiaga RD, Stets CM, Katzenstein HM, Millis SZ, Rogers A, et al. Multiscale-omic assessment of EWSR1-NFATc2 fusion positive sarcomas identifies the mTOR pathway as a potential therapeutic target. NPJ Precis Oncol. 2021;5:43.

18. Mateo-Lozano S, Tirado OM, Notario V. Rapamycin induces the fusion-type independent downregulation of the EWS/FLI-1 proteins and inhibits Ewing’s sarcoma cell proliferation. Oncogene. Nature Publishing Group; 2003;22:9282–7.

19. Zenali MJ, Zhang PL, Bendel AE, Brown RE. Morphoproteomic Confirmation of Constitutively Activated mTOR, ERK, and NF-kappaB Pathways in Ewing Family of Tumors. Ann Clin Lab Sci. Association of Clinical Scientists; 2009;39:160–6.

20. Guenther LM, Dharia NV, Ross L, Conway A, Robichaud AL, Catlett JL, et al. A Combination CDK4/6 and IGF1R Inhibitor Strategy for Ewing Sarcoma. Clin Cancer Res Off J Am Assoc Cancer Res. 2019;25:1343– 57.

21. Crompton BD, Carlton AL, Thorner AR, Christie AL, Du J, Calicchio ML, et al. High-Throughput Tyrosine Kinase Activity Profiling Identifies FAK as a Candidate Therapeutic Target in Ewing Sarcoma. Cancer Res. 2013;73:2873–83.

22. Kennedy AL, Vallurupalli M, Chen L, Crompton B, Cowley G, Vazquez F, et al. Functional, chemical genomic, and super-enhancer screening identify sensitivity to cyclin D1/CDK4 pathway inhibition in Ewing sarcoma. Oncotarget. Impact Journals; 2015;6:30178–93.

23. Wakahara K, Ohno T, Kimura M, Masuda T, Nozawa S, Dohjima T, et al. EWS-Fli1 Up-Regulates Expression of the Aurora A and Aurora B Kinases. Mol Cancer Res. 2008;6:1937–45.

24. Potratz JC, Saunders DN, Wai DH, Ng TL, McKinney SE, Carboni JM, et al. Synthetic Lethality Screens Reveal RPS6 and MST1R as Modifiers of Insulin-like Growth Factor-1 Receptor Inhibitor Activity in Childhood Sarcomas. Cancer Res. 2010;70:8770–81.

25. Wang S, Hwang EE, Guha R, O’Neill AF, Melong N, Veinotte CJ, et al. High-throughput Chemical Screening Identifies Focal Adhesion Kinase and Aurora Kinase B Inhibition as a Synergistic Treatment Combination in Ewing Sarcoma. Clin Cancer Res. 2019;25:4552–66.

26. Mathews Griner LA, Guha R, Shinn P, Young RM, Keller JM, Liu D, et al. High-throughput combinatorial screening identifies drugs that cooperate with ibrutinib to kill activated B-cell–like diffuse large B-cell lymphoma cells. Proc Natl Acad Sci U S A. 2014;111:2349–54.

27. Tibes R, Qiu Y, Lu Y, Hennessy B, Andreeff M, Mills GB, et al. Reverse phase protein array: validation of a novel proteomic technology and utility for analysis of primary leukemia specimens and hematopoietic stem cells. Mol Cancer Ther. 2006;5:2512–21.

28. Davis JB, Andes S, Espina V. Reverse Phase Protein Arrays. Methods Mol Biol Clifton NJ. 2021;2237:103– 22.

29. Zhu A, Ibrahim JG, Love MI. Heavy-tailed prior distributions for sequence count data: removing the noise and preserving large differences. Bioinformatics. 2019;35:2084–92.

30. Vora SR, Juric D, Kim N, Mino-Kenudson M, Huynh T, Costa C, et al. CDK 4/6 Inhibitors Sensitize PIK3CA Mutant Breast Cancer to PI3K Inhibitors. Cancer Cell. 2014;26:136–49.

31. Gaspar N, Hawkins DS, Dirksen U, Lewis IJ, Ferrari S, Le Deley M-C, et al. Ewing Sarcoma: Current Management and Future Approaches Through Collaboration. J Clin Oncol Off J Am Soc Clin Oncol. 2015;33:3036–46.

32. Povedano JM, Li V, Lake KE, Bai X, Rallabandi R, Kim J, et al. TK216 targets microtubules in Ewing sarcoma cells. Cell Chem Biol. 2022;29:1325–1332.e4.

33. Erkizan HV, Kong Y, Merchant M, Schlottmann S, Barber-Rotenberg JS, Abaan OD, et al. Small molecule selected to disrupt oncogenic protein EWS-FLI1 interaction with RNA Helicase A inhibits Ewing’s Sarcoma. Nat Med. 2009;15:750–6.

34. Barber-Rotenberg JS, Selvanathan SP, Kong Y, Erkizan HV, Snyder TM, Hong SP, et al. Single Enantiomer of YK-4-279 Demonstrates Specificity in Targeting the Oncogene EWS-FLI1. Oncotarget. 2012;3:172–82.

35. Meyers PA, Federman N, Daw N, Anderson Peter M, Davis LE, Kim A, et al. Open-Label, Multicenter, Phase I/II, First-in-Human Trial of TK216: A First-Generation EWS::FLI1 Fusion Protein Antagonist in Ewing Sarcoma. J Clin Oncol. Wolters Kluwer; 2024;0:JCO.24.00020.

36. Theisen ER, Selich-Anderson J, Miller KR, Tanner JM, Taslim C, Pishas KI, et al. Chromatin profiling reveals relocalization of lysine-specific demethylase 1 by an oncogenic fusion protein. Epigenetics. 16:405–24.

37. Sankar S, Theisen ER, Bearss J, Mulvihill T, Hoffman LM, Sorna V, et al. Reversible LSD1 Inhibition Interferes with Global EWS/ETS Transcriptional Activity and Impedes Ewing Sarcoma Tumor Growth. Clin Cancer Res. 2014;20:4584–97.

38. Reed DR, Chawla SP, Setty B, Mascarenhas L, Meyers PA, Metts J, et al. Phase 1 trial of seclidemstat (SP- 2577) in patients with relapsed/refractory Ewing sarcoma. J Clin Oncol. Wolters Kluwer; 2021;39:11514– 11514.

39. Grohar PJ, Ballman KV, Heise R, Mascarenhas L, Glod J, Wedekind MFF, et al. SARC037: Phase II results of trabectedin given as a 1-hour (h) infusion in combination with low dose irinotecan in patients (pts) with relapsed/refractory Ewing sarcoma (ES). J Clin Oncol. Wolters Kluwer; 2024;42:11508–11508.

40. Harlow ML, Maloney N, Roland J, Guillen Navarro MJ, Easton MK, Kitchen-Goosen SM, et al. Lurbinectedin inactivates the Ewing sarcoma oncoprotein EWS-FLI1 by redistributing it within the nucleus. Cancer Res. 2016;76:6657–68.

41. Harlow ML, Chasse MH, Boguslawski EA, Sorensen KM, Gedminas JM, Kitchen-Goosen SM, et al. Trabectedin Inhibits EWS-FLI1 and Evicts SWI/SNF from Chromatin in a Schedule Dependent Manner. Clin Cancer Res Off J Am Assoc Cancer Res. 2019;25:3417–29.

42. Grohar PJ, Segars LE, Yeung C, Pommier Y, D’Incalci M, Mendoza A, et al. Dual targeting of EWS-FLI1 activity and the associated DNA damage response with Trabectedin and SN38 synergistically inhibits Ewing sarcoma cell growth. Clin Cancer Res Off J Am Assoc Cancer Res. 2014;20:1190–203.

43. Pappo AS, Patel SR, Crowley J, Reinke DK, Kuenkele K-P, Chawla SP, et al. R1507, a Monoclonal Antibody to the Insulin-Like Growth Factor 1 Receptor, in Patients With Recurrent or Refractory Ewing Sarcoma Family of Tumors: Results of a Phase II Sarcoma Alliance for Research Through Collaboration Study. J Clin Oncol. 2011;29:4541–7.

44. Shulman DS, Merriam P, Choy E, Guenther LM, Cavanaugh KL, Kao P, et al. Phase 2 trial of palbociclib and ganitumab in patients with relapsed Ewing sarcoma. Cancer Med. 2023;12:15207–16.

45. Dharia NV, Kugener G, Guenther LM, Malone CF, Durbin AD, Hong AL, et al. A First-Generation Pediatric Cancer Dependency Map. Nat Genet. 2021;53:529–38.

