## Supplemental Figures for "A novel combination of CDK4/6 and PI3K inhibitors exhibits highly synergistic activity and translational potential in Ewing sarcoma"

### Supplemental Figure 1

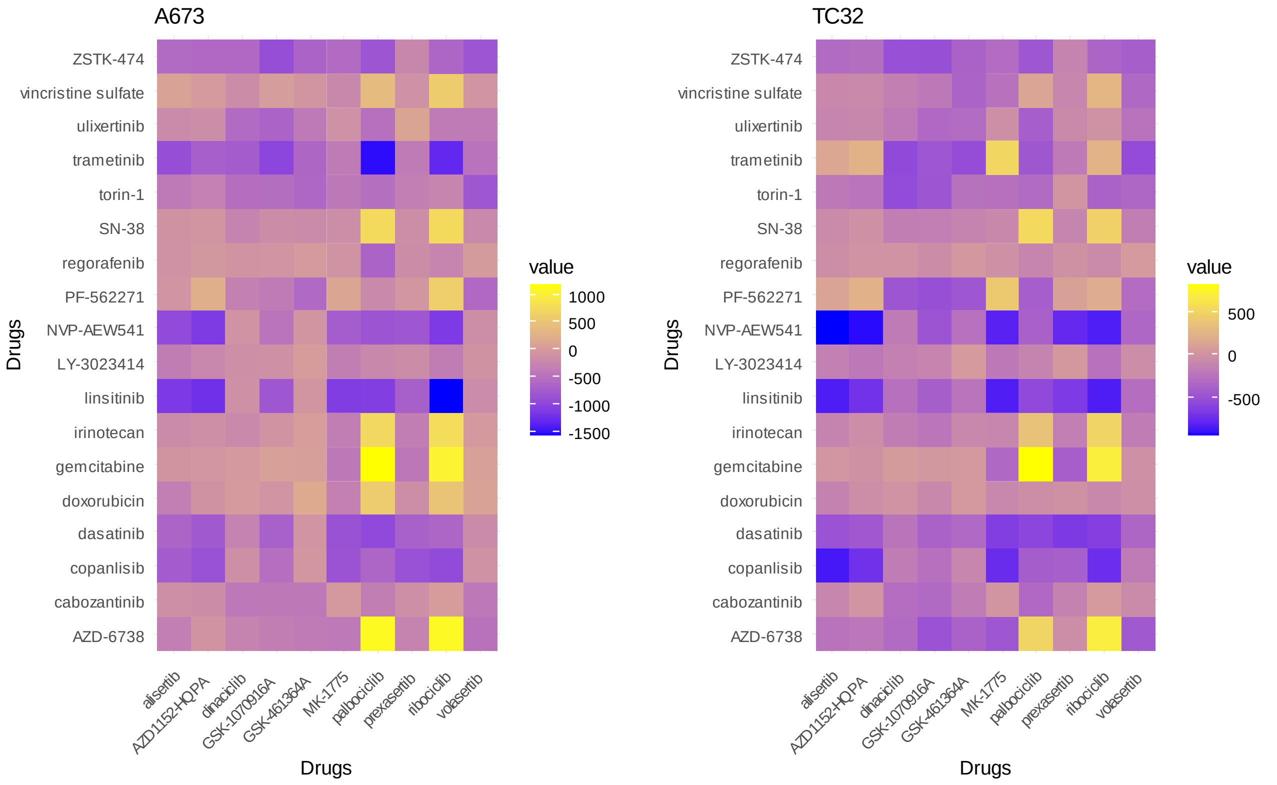

### Supplemental Figure 2

## A

#### A673 Ewing cells

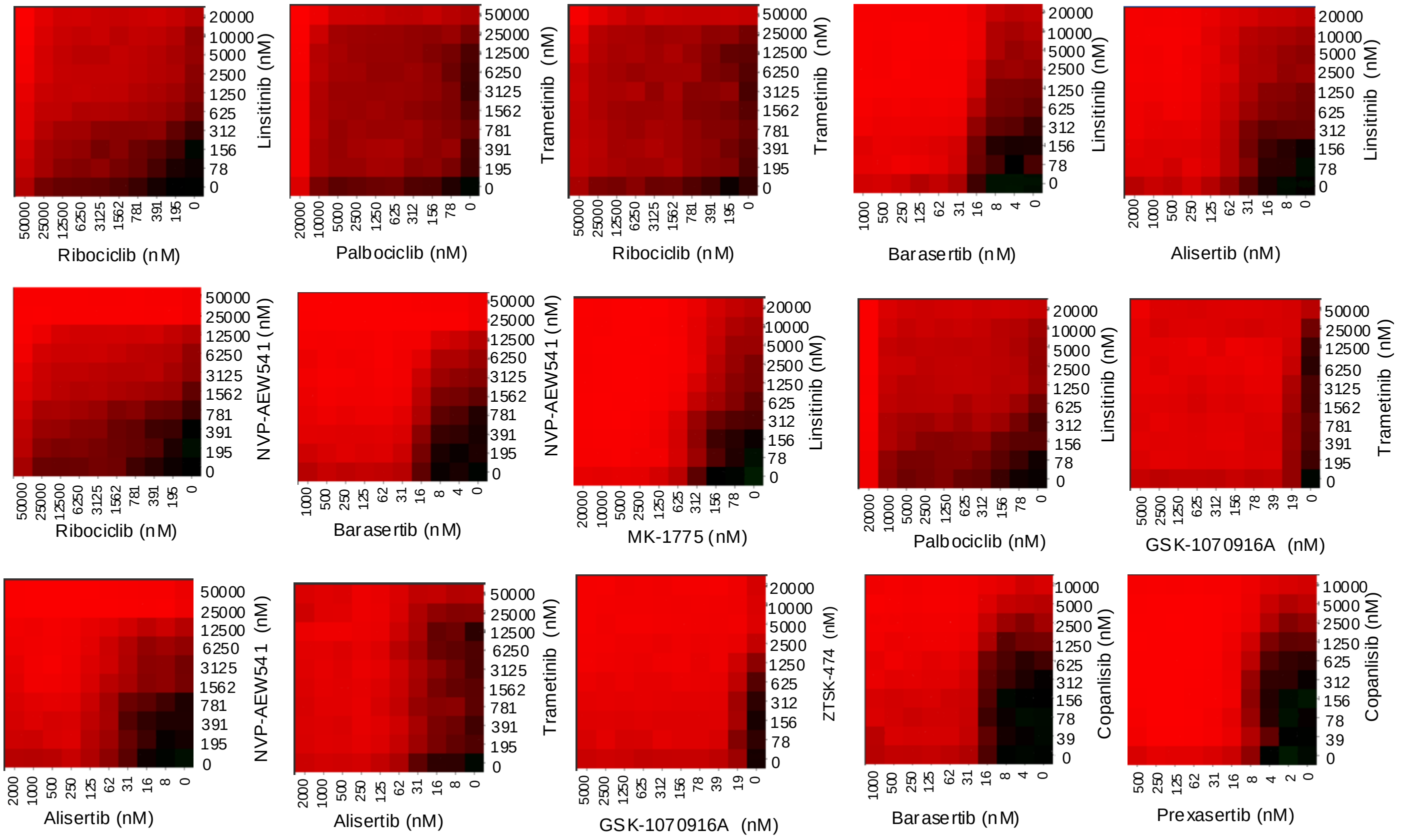


## B

#### TC32 Ewing cells

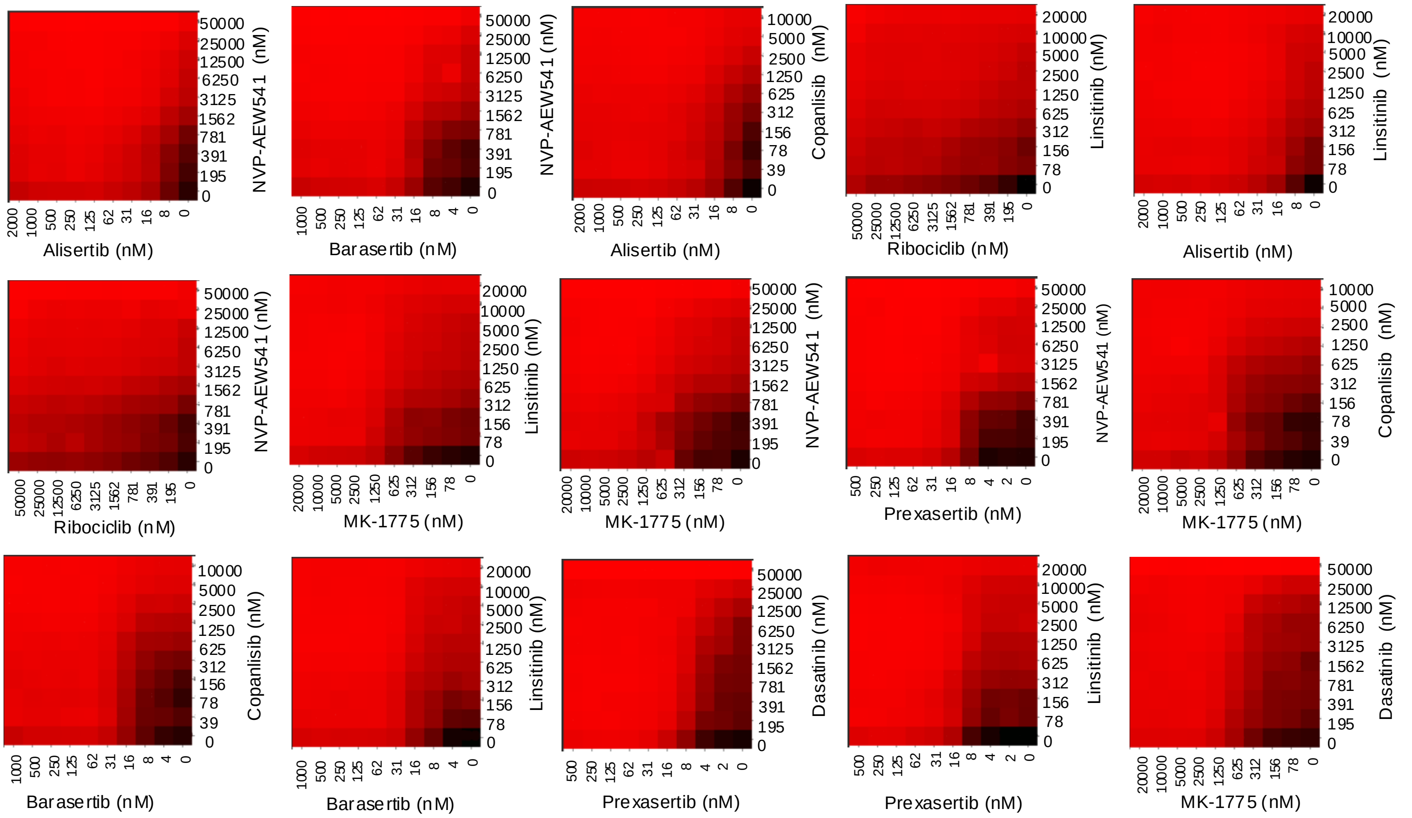

### Supplemental Figure 3

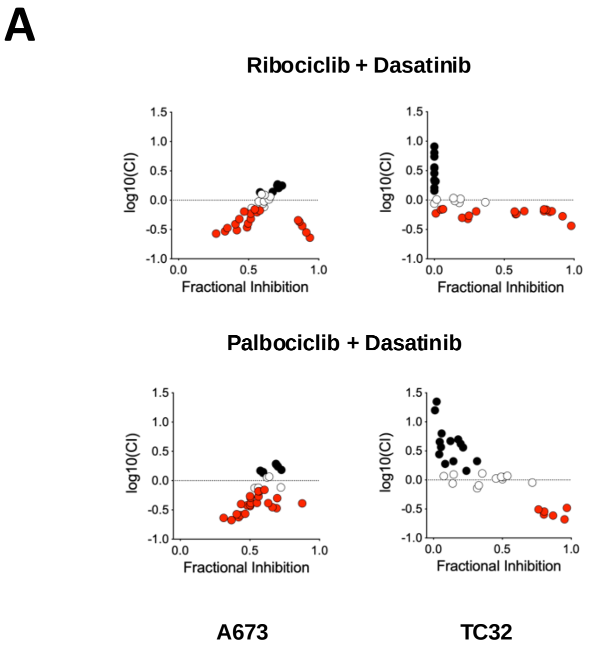


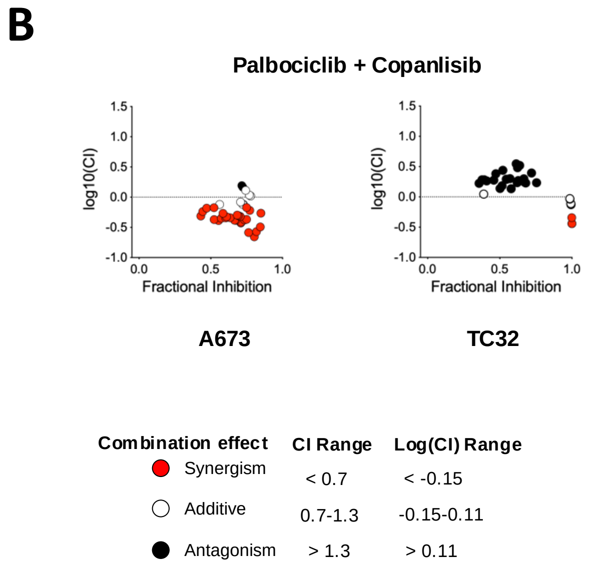

### Supplemental Figure 4

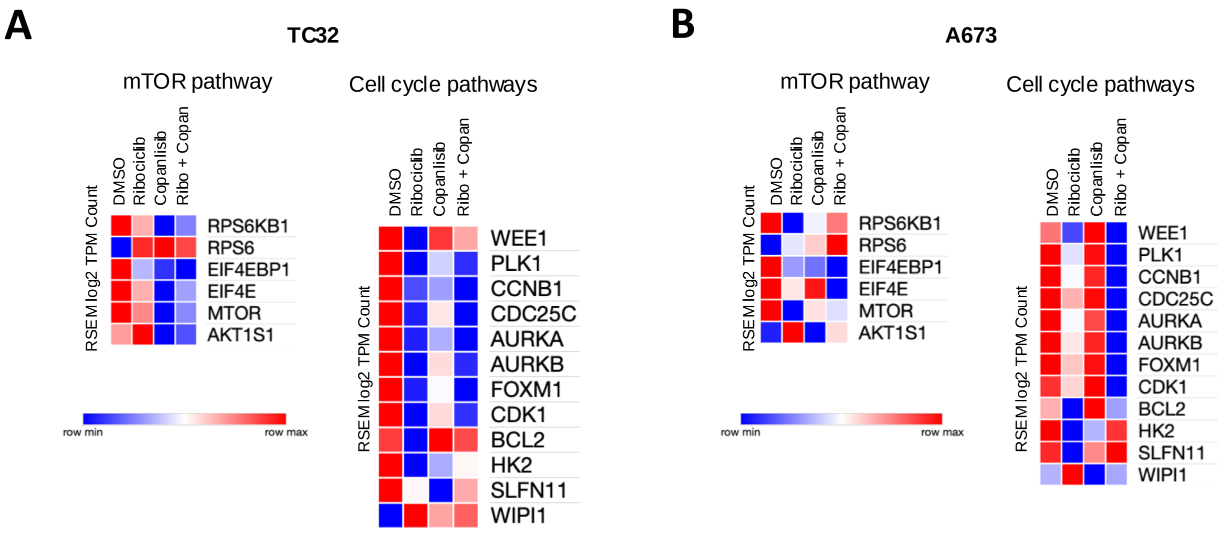

### Supplemental Figure 5

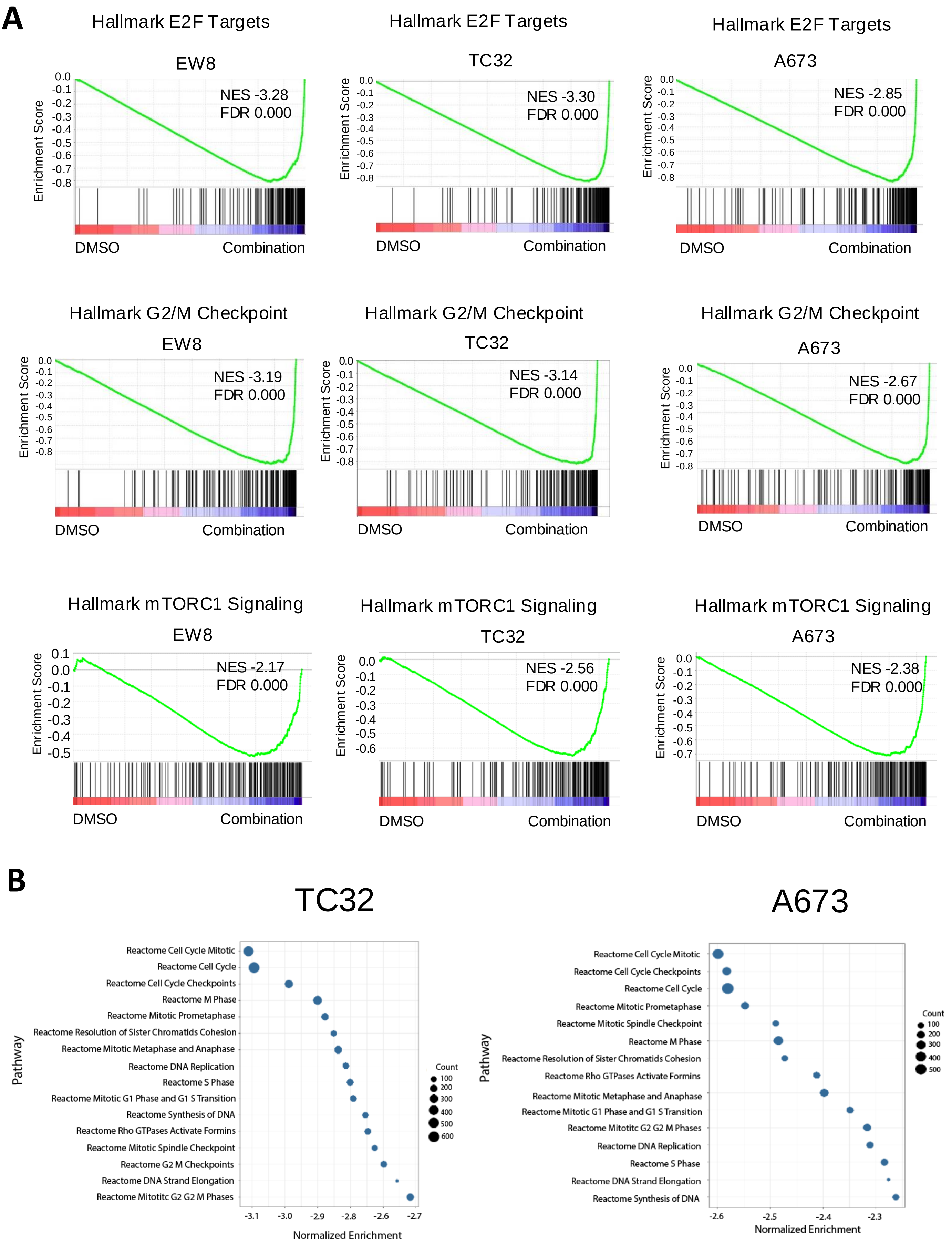

### Supplemental Figure 6

#### Efficacy cohort

A

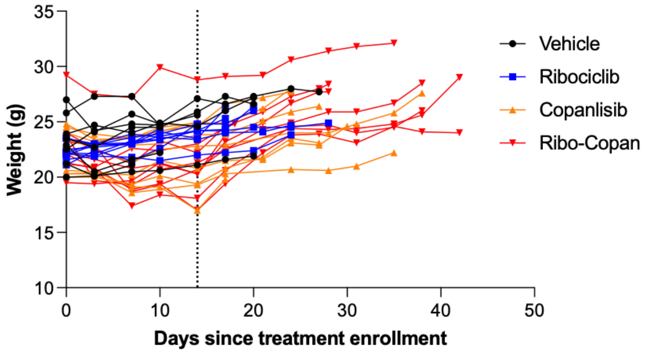


B

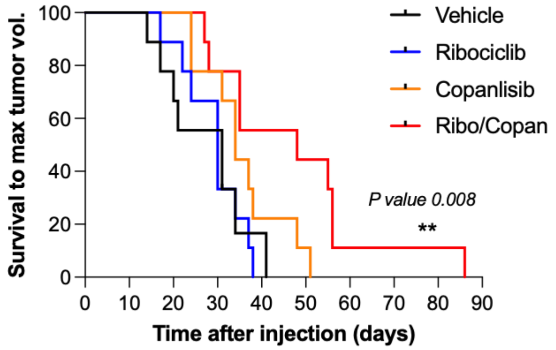


C

#### Survival cohort

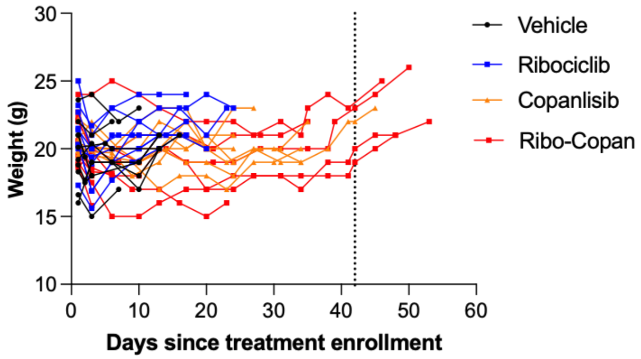
