## Supplemental Tables for "A novel combination of CDK4/6 and PI3K inhibitors exhibits highly synergistic activity and translational potential in Ewing sarcoma"

| Drug | Mechanism | Category | IC50 Range A673 | IC50 Range TC32 | Starting dose | Dilution factor |
|---|---|---|---|---|---|---|
| Ulixertinib | ERK1/2 Inhibitor | Receptor tyrosine kinase inhibitor | 1.46-16.53 uM | 0.37-2.42 uM | 20 uM | 1:2 dilution |
| Cabozantinib | VEGFR-2 Inhibitor | Receptor tyrosine kinase inhibitor | 6.09-12.6 uM | 5.68-11.7 uM | 20 uM | 1:2 dilution |
| Trametinib | MEK 1/2 Inhibitor | Receptor tyrosine kinase inhibitor | 3.68-55 uM | 2 uM | 20 uM | 1:2 dilution |
| Regorafenib | VEGFR-2 Inhibitor | Receptor tyrosine kinase inhibitor | 10-16.54 uM | 1.62-8.87 uM | 20 uM | 1:2 dilution |
| Linsitinib | IGF-1R inhibitor | Receptor tyrosine kinase inhibitor | 0.4-11.16 uM | 0.1-3.8 uM | 20 uM | 3:4 dilution |
| NVP-AEW541 | IGF-1R inhibitor | Receptor tyrosine kinase inhibitor | 4.13-18.55 uM | 0.92-16 uM | 20 uM | 3:4 dilution |
| Dasatinib | Bcr-Abl Inhibitor | Receptor tyrosine kinase inhibitor | 2.5-17.6 uM | N/A | 20 uM | 1:2 dilution |
| LY-3023414 | PI3K Inhibitor | Kinase inhibitor | 0.15-0.47 uM | 0.11-0.37 uM | 2 uM | 1:2 dilution |
| Copanlisib | PI3K alpha/delta inhibitor | Kinase inhibitor | 1.16-2.8 uM | 0.06-0.42 uM | 1-5 uM | 1:2 dilution |
| ZSTK-474 | PI3K Inhibitor | Kinase inhibitor | 0.77-2.62 uM | 0.24-0.54 uM | 20 uM | 1:2 dilution |
| PF-562271 | FAK inhibitor | Kinase inhibitor | 2.6-6.6 uM | 2.5-4.7 uM | 10-20 uM | 1:2 dilution |
| Torin-1 | mTOR inhibitor | mTOR pathway | 51.5-203 nM | 30.5-220 nM | 5 uM | 1:2 dilution |
| Prexasertib | Chk1 inhibitor | Cell cycle inhibitor | 3.7-10.25 nM | 1.5-8.3 | 100 nM | 1:2 dilution |
| Volasertib | Plk-1 inhibitor | Cell cycle inhibitor | 15.56-83 nM | 29.9-80.2 nM | 0.5 uM | 1:2 dilution |
| Dinaciclib | CDK1/2/5/9 | Cell cycle inhibitor | 9.5-25 nM | 7.8-23 nM | 100 nM | 1:2 dilution |
| Alisertib | AURKA inhibitor | Cell cycle inhibitor | 34.4 nM-1.46 uM | 8.9-46 nM | 1-5 uM | 1:2 dilution |
| GSK-461364A | Plk-1 inhibitor | Cell cycle inhibitor | 52.6-106.4 | 60.2 nM | 1 uM | 1:2 dilution |
| Adavosertib/MK1775 | Wee1 Kinase inhibitor | Cell cycle inhibitor | 0.28-0.46 uM | 0.43-0.83 uM | 5 uM | 1:2 dilution |
| Palbociclib | CDK4/6 inhibitor | Cell cycle inhibitor | 2.24-3.7 uM | 1-2 uM | 20 uM | 1:2 dilution |
| Barasertib | AURKB inhibitor | Cell cycle inhibitor | 15-25 nM | 10-25 nM | 200 nM | 1:2 dilution |
| Ribociclib | CDK4/6 inhibitor | Cell cycle inhibitor | 2.4-10.43 uM | 3.8-5.87 uM | 20 uM | 3:4 dilution |
| GSK-1070916 | AURKB/C inhibitor | Cell cycle inhibitor | 14-16 nM | 23-93 nM | 100-500 nM | 1:2 dilution |
| Gemcitabine | Ribonucleotide reductase inhibitor | Chemotherapy | 3.47-11.02 nM | 3.04-6.8 nM | 20 nM | 1:2 dilution |
| Vincristine Sulfate | Tubulin polymerization inhibitor | Chemotherapy | 1.67-6 nM | 2.4-2.77 nM | 20 nM | 1:2 dilution |
| SN-38 | DNA Topoisomerase 1 Inhibitor | Chemotherapy | 1.64-2.13 nM | 0.34-1.9 nM | 10 nM | 1:2 dilution |
| Irinotecan | DNA Topoisomerase 1 Inhibitor | Chemotherapy | 61.9-65.8 nM | 24.1-131 nM | 1 uM | 1:2 dilution |
| Doxorubicin | DNA Topoisomerase II Inhibitor | Chemotherapy | 25-66 uM | 2.25-13 nM | 0.5 uM | 1:2 dilution |
| AZD-6738 | ATR Kinase inhibitor | ATR inhibitor | 0.69-5.22 uM | 0.42-2.2 uM | 10-20 uM | 1:2 dilution |

**Supplemental Table 1: Single-agent dose response curve from the 28 selected compounds.**

| A673 | | | | | |
|---|---|---|---|---|---|
| Ranking of top 10 percent combinations | Drug 1 | Drug 1 description | Drug 2 | Drug 2 description | Excess HSA |
| 1 | Linsitinib | IGF-1R Inhibitor | Ribociclib (LEE011) | CDK4/6 Inhibitor | -1556.9553 |
| 2 | Trametinib | Mek 1/2 Inhibitor | Palbociclib | CDK4/6 Inhibitor | -1501.6418 |
| 3 | Trametinib | Mek 1/2 Inhibitor | Ribociclib (LEE011) | CDK4/6 Inhibitor | -1293.6345 |
| 4 | Linsitinib | IGF-1R Inhibitor | Barasertib (AZD1152-HQPA) | Aurora-B Inhibitor | -1212.2062 |
| 5 | Linsitinib | IGF-1R Inhibitor | Alisertib | Aurora-A Inhibitor | -1124.9619 |
| 6 | NVP-AEW541 | Unknown | Ribociclib (LEE011) | CDK4/6 Inhibitor | -1109.335 |
| 7 | NVP-AEW541 | Unknown | Barasertib (AZD1152-HQPA) | Aurora-B Inhibitor | -1105.556 |
| 8 | Linsitinib | IGF-1R Inhibitor | MK-1775 | Wee1 Kinase Inhibitor | -1074.5439 |
| 9 | Linsitinib | IGF-1R Inhibitor | Palbociclib | CDK4/6 Inhibitor | -1066.8058 |
| 10 | Trametinib | Mek 1/2 Inhibitor | GSK-1070916A | Aurora-B/C Inhibitor | -986.3008 |
| 11 | Dasatinib | Bcr-Abl Inhibitor | Palbociclib | CDK4/6 Inhibitor | -951.62125 |
| 12 | NVP-AEW541 | Unknown | Alisertib | Aurora-A Inhibitor | -940.88245 |
| 13 | Copanlisib | PI3K alpha/delta Inhibitor | Ribociclib (LEE011) | CDK4/6 Inhibitor | -928.4249 |
| 14 | Trametinib | Mek 1/2 Inhibitor | Alisertib | Aurora-A Inhibitor | -895.29946 |
| 15 | ZSTK-474 | PI3K Inhibitor | GSK-1070916A | Aurora-B/C Inhibitor | -893.01892 |
| 16 | Dasatinib | Bcr-Abl Inhibitor | Ribociclib (LEE011) | CDK4/6 Inhibitor | -886.87707 |
| 17 | Copanlisib | PI3K alpha/delta Inhibitor | Barasertib (AZD1152-HQPA) | Aurora-B Inhibitor | -861.26294 |
| 18 | Copanlisib | PI3K alpha/delta Inhibitor | Prexasertib | Chk1 Inhibitor | -858.84589 |

**Supplemental Table 2: Top 10 percent excess over the highest single drug response (HAS) in A673 Ewing cells**

| TC32 | | | | | |
|---|---|---|---|---|---|
| **Ranking of top 10 percent combinations** | **Drug 1** | **Drug 1 description** | **Drug 2** | **Drug 2 description** | **Excess HSA** |
| 1 | NVP-AEW541 | Unknown | Alisertib | Aurora-A Inhibitor | -948.49469 |
| 2 | NVP-AEW541 | Unknown | Barasertib (AZD1152-HQPA) | Unknown | -918.38006 |
| 3 | Copanlisib | PI3K alpha/delta Inhibitor | Alisertib | Aurora-A Inhibitor | -863.05045 |
| 4 | Linsitinib | IGF-1R Inhibitor | Ribociclib (LEE011) | CDK4/6 Inhibitor | -844.80906 |
| 5 | Linsitinib | IGF-1R Inhibitor | Alisertib | Aurora-A Inhibitor | -844.22159 |
| 6 | NVP-AEW541 | Unknown | Ribociclib (LEE011) | CDK4/6 Inhibitor | -840.64166 |
| 7 | Linsitinib | IGF-1R Inhibitor | MK-1775 | Wee1 Kinase Inhibitor | -838.85888 |
| 8 | NVP-AEW541 | Unknown | MK-1775 | Wee1 Kinase Inhibitor | -812.18038 |
| 9 | NVP-AEW541 | Unknown | Prexasertib | Chk1 Inhibitor | -774.39314 |
| 10 | Copanlisib | PI3K alpha/delta Inhibitor | MK-1775 | Wee1 Kinase Inhibitor | -751.9052 |
| 11 | Copanlisib | PI3K alpha/delta Inhibitor | Ribociclib (LEE011) | CDK4/6 Inhibitor | -742.7622 |
| 12 | Copanlisib | PI3K alpha/delta Inhibitor | Barasertib (AZD1152-HQPA) | Aurora-B Inhibitor | -719.10344 |
| 13 | Linsitinib | IGF-1R Inhibitor | Barasertib (AZD1152-HQPA) | Aurora-B Inhibitor | -714.45353 |
| 14 | Dasatinib | Bcr-Abl Inhibitor | Prexasertib | Chk1 Inhibitor | -659.90092 |
| 15 | Linsitinib | IGF-1R Inhibitor | Prexasertib | Chk1 Inhibitor | -656.99806 |
| 16 | Dasatinib | Bcr-Abl Inhibitor | MK-1775 | Wee1 Kinase Inhibitor | -629.25032 |
| 17 | Dasatinib | Bcr-Abl Inhibitor | Ribociclib (LEE011) | CDK4/6 Inhibitor | -621.76764 |
| 18 | Dasatinib | Bcr-Abl Inhibitor | Palbociclib | CDK4/6 Inhibitor | -580.20017 |

**Supplemental Table 3: Top 10 percent excess over the highest single drug response (HAS) in TC32 Ewing cells**

| Efficacy cohort - Time to reach endpoint (days) | | | | | | | |
|---|---|---|---|---|---|---|---|
| **Mouse ID** | **Vehicle** | **Mouse ID** | **Ribociclib** | **Mouse ID** | **Copanlisib** | **Mouse ID** | **Combination** |
| C1_none | 14.0 | C1_none | 10.0 | C1_none | 24.0 | C1_none | 28.0 |
| C1_L | 7.0 | C1_L | 17.0 | C1_L | 24.0 | C1_L | 18.0 |
| C1_R | 10.0 | C1_R | 14.0 | C1_R | 30.0 | C1_R | 45.0 |
| C1_none | 10.0 | C1_none | 21.0 | C1_none | 17.0 | C1_none | 28.0 |
| C2_L | 21.0 | C2_L | 22.0 | C2_L | 24.0 | C2_L | 38.0 |
| C2_R | 31.0 | C2_R | 19.0 | C2_R | 14.0 | C2_R | 18.0 |
| C2_LR | 21.0 | C2_LR | 28.0 | C2_LR | 28.0 | C2_LR | 43.0 |
| C2_none | 10 | C2_none | 24 | C2_none | 38.0 | C2_none | 60.0 |
| C2_R | 17 | C2_R | 17 | C2_R | 42.0 | C2_R | 43 |
| Average (days) | 15.7 | | 19.1 | | 26.8 | | 35.7 |

**Supplemental Table 4: Time to reach endpoint after beginning of therapy in the efficacy cohort.**

| Efficacy cohort - Tumor size (mm3) | | | | | | | |
|---|---|---|---|---|---|---|---|
| **Mouse ID** | **Vehicle** | **Mouse ID** | **Ribociclib** | **Mouse ID** | **Copanlisib** | **Mouse ID** | **Combination** |
| C1_none | 657.0 | C1_none | 1007.0 | C1_none | 151.0 | C1_none | 127.0 |
| C1_L | 1942.0 | C1_L | 595.0 | C1_L | 126.0 | C1_L | 262.0 |
| C1_R | 1328.0 | C1_R | 779.0 | C1_R | 137.0 | C1_R | 59.0 |
| C1_none | 789.0 | C1_none | 436.0 | C1_none | 454.0 | C1_none | 52.0 |
| C2_L | 231.0 | C2_L | 442.0 | C2_L | 146.0 | C2_L | 78.0 |
| C2_R | 117.0 | C2_R | 295.0 | C2_R | 1021.0 | C2_R | 605.0 |
| C2_LR | 298.0 | C2_LR | 111.0 | C2_LR | 301.0 | C2_LR | 32.0 |
| C2_none | 1293 | C2_none | 431 | C2_none | 92.0 | C2_none | 78.0 |
| C2_R | 116 | C2_R | 262 | C2_R | 128.0 | C2_R | 32 |
| Average (mm3) | 752.3 | | 484.2 | | 284.0 | | 161.6 |

**Supplemental Table 5: Tumor size 7 days after enrollment in the efficacy cohort.**

| Survival cohort (Days post injection) | | | | | | | | | | | |
|---|---|---|---|---|---|---|---|---|---|---|---|
| Mouse ID | Vehicle | Toxicity | Mouse ID | Ribociclib | Toxicity | Mouse ID | Copanlisib | Toxicity | Mouse ID | Combination | Toxicity |
| C1_none | 27.0 | | C1_none | 30.0 | | C1_none | 39.0 | | C1_none* | 33.0 | weight loss |
| C1_L* | 14.0 | oral gavage | C1_L | 29.0 | | C1_L | 35.0 | | C1_L | 60.0 | |
| C1_R | 24.0 | | C1_R* | 30.0 | oral gavage | C1_R | 43.0 | | C1_R* | 41.0 | weight loss |
| C1_LR | 30.0 | | C1_LR | 35.0 | | C1_2L* | 43.0 | GI distress | C1_LR* | 13.0 | oral gavage |
| C1_2L | 24.0 | | C1_2L | 32.0 | | C2_none | 57.0 | | C1_2R | 70.0 | |
| C2_none | 28.0 | | C2_none | 35.0 | | C2_L | 32.0 | | C2_none* | 17.0 | oral gavage |
| C2_L | 28.0 | | C2_L | 27.0 | | C2_R | 49.0 | | C2_L | 67.0 | |
| C2_R* | 17.0 | oral gavage | C2_R* | 17.0 | oral gavage | C2_LR | 38.0 | | C2_LR | 52 | |
| C2_LR | 24 | | C2_LR | 32 | | | | | C2_2L | 60 | |
| C2_2L | 21 | | | | | | | | | | |
| Average (days) | 25.8 | | | 31.4 | | | 42.0 | | | 60.0 | |

**Supplemental Table 6: Time to reach endpoint after subcutaneous injection in the survival cohort.**

* weight loss: animal censored due weight loss greater than 10% of their original weight at enrollment
* oral gavage: animal censored after necropsy revealed evidence of esophageal perforation due to intense daily schedule of oral gavage procedure for 6 weeks
* toxicity: animal censored after necropsy revealed signs of gastrointestinal distress or liver dysfunction which could be signs of treatment toxicity

| Study title | NCT number | Status | Conditions | Intervention / Treatment | Study Type |
|---|---|---|---|---|---|
| Testing the safety and efficacy of the combination of two anti-cancer drugs, ZEN003694 and Abemaciclib, for Adult and Pediatric Patients (12-17 years) with metastatic or unresectable breast cancer and other solid tumors | NCT05372640 | Recruiting | Breast carcinoma<br>Malignant solid neoplasm<br>NUT Carcinoma | • Drug: Abemaciclib<br>• Drug: BET Bromodomain inhibitor ZEN-3694 | Phase 1 |
| Study of Palbociclib Combined With Chemotherapy in Pediatric Patients With Recurrent/Refractory Solid Tumors | NCT03709680 | Active, not recruiting | Ewing Sarcoma<br>Solid Tumors<br>Rhabdoid Tumor<br>Neuroblastoma<br>Medulloblastoma<br>Diffuse Intrinsic Pontine Glioma | • Drug: Palbociclib<br>• Drug: Temozolomide<br>• Drug: Irinotecan<br>• Drug: Topotecan<br>• Drug: Cyclophosphamide | Phase 1/2 |
| Abemaciclib in Children with DIPG or Recurrent/Refractory Solid Tumors | NCT02644460 | Completed | Diffuse Intrinsic Pontine Glioma<br>Brain Tumor, Recurrent<br>Neuroblastoma, Recurrent, Refractory<br>Ewing Sarcoma, Recurrent, Refractory<br>Rhadomyosarcoma, Recurrent, Refractory<br>Osteosarcoma, Recurrent, Refractory<br>Rhabdoid Tumor, Recurrent, Refractory | • Drug: Abemaciclib | Phase 1 |
| Safety, Tolerability, Efficacy, and Pharmacokinetics of Copanlisib in Pediatric patients | NCT03458728 | Completed | Relapsed or Refractory Solid Tumors or Lymphoma in Children<br>Neuroblastoma<br>Osteosarcoma<br>Rhadomyosarcoma<br>Ewing sarcoma | • Drug: Copanlisib | Phase 1/2 |
| Palbociclib + Ganitumab in Ewing Sarcoma | NCT04129151 | Completed | Ewing Sarcoma | • Drug: Palbociclib<br>• Drug: Ganitumab | Phase 1/2 |
| Study of Safety and Efficacy in Patients with Malignant Rhabdoid Tumors (MRT) and Neuroblastoma | NCT01747876 | Completed | Malignant Rhabdoid Tumors, Neuroblastoma | • Drug: Ribociclib (LEE011) | Phase 1 |
| Palbociclib Isethionate in Treating Younger Patients With Recurrent, Progressive, or Refractory Central Nervous System Tumors | NCT02255461 | Terminated | Childhood Choroid Plexus Tumor, Childhood Ependymoblastoma, Childhood Grade III Meningioma, Childhood High-grade Cerebellar Astrocytoma, Childhood High-grade Cerebral Astrocytoma, Childhood Medulloepithelioma, Recurrent Childhood Anaplastic Astrocytoma, Recurrent Childhood Brain Stem Glioma, Recurrent Childhood Cerebellar Astrocytoma, Recurrent Childhood Cerebral Astrocytoma, Recurrent Childhood Giant Cell Glioblastoma, Recurrent Childhood Glioblastoma, Recurrent Childhood Gliomatosis Cerebri, Recurrent Childhood Supratentorial Primitive Neuroectodermal Tumor | • Drug: Palbociclib isethionate | Phase 1 |
| St Jude Children's Research Hospital Phase 1 Study Evaluating Molecularly-Driven Doublet Therapies for Children and Young Adults with Recurrent Brain Tumors | NCT03434262 | Completed | Recurrent or relasped malignant brain tumors | • Drug: Gemcitabine<br>• Drug: Ribociclib<br>• Drug: Sonidegib<br>• Drug: Trametinib | Phase 1 |
| Study of efficacy and safety of Ribociclib (LEEO11) in combination with Topotecan and Temozolomide (TOTEM) in Pediatric patients with relapsed or refractory neuroblastoma and other solid tumors | NCT05429502 | Recruiting | Neuroblastoma | • Drug: Topotecan<br>• Drug: Temozolomide<br>• Drug: Ribociclib | Phase I-part A (dose finding) and Phase I-part B (multiple expansion cohorts) |

**Table 7: Summary of clinical trials involving the use of PI3K or CDK4/6 inhibitors.**
* Source: clinicaltrials.gov
